# VEGFR2 blockade overcomes acquired KRAS G12D inhibitor resistance driven by PI3Kγ activation

**DOI:** 10.1101/2025.09.16.676456

**Authors:** Sung-Hyun Hwang, Mingyun Bae, Ji-Won Kim, Seung Yoon Hyun, Kui-Jin Kim, Joshua H. Choe, Min Jung Kim, Ji Won Park, Seung-Yong Jeong, Songji Choi, Woochan Park, Jeongmin Seo, Heejung Chae, Minsu Kang, Eun Hee Jung, Koung Jin Suh, Se Hyun Kim, Jin Won Kim, Yu Jung Kim, Jee Hyun Kim, Haeseong Park, Andrew J. Aguirre, Eunjung Alice Lee, Ja-Lok Ku, Keun-Wook Lee

**Affiliations:** Biomedical Research Institute, Seoul National University Bundang Hospital, Seongnam 13620, Republic of Korea; Division of Genetics and Genomics, Boston Children’s Hospital, Harvard Medical School, Boston, MA 02115, United States; Broad Institute of MIT and Harvard, Cambridge, MA 02142, USA; Department of Internal Medicine, Seoul National University Bundang Hospital, Seoul National University College of Medicine, Seongnam 13620, Republic of Korea; Department of Genomic Medicine, Seoul National University Bundang Hospital, Seongnam 13620, Republic of Korea; Laboratory of Cell Biology, Cancer Research Institute, Seoul National University College of Medicine, Seoul 03080, Republic of Korea; Department of Medical Oncology, Dana-Farber Cancer Institute, Boston, MA 02215, USA; Division of Medical Sciences, Harvard Medical School, Boston, MA 02215; Department of Surgery, Seoul National University Hospital, Seoul National University College of Medicine, Seoul 03080, Republic of Korea; Department of Medicine, Brigham and Women’s Hospital and Harvard Medical School, Boston, MA 02115, USA

**Keywords:** KRAS, MRTX1133, acquired resistance, angiogenesis, VEGFR2

## Abstract

KRAS G12D mutation is a key oncogenic driver in many solid tumors, including pancreatic, gastric, and colorectal cancers. While recent studies have characterized features associated with primary and acquired resistance to KRAS inhibitors, strategies to overcome such resistance, particularly in the context of gastrointestinal cancers, remain underexplored. Here, we have generated nine human gastrointestinal cancer models, including three patient-derived organoids (PDOs), with acquired resistance to the KRAS G12D-selective inhibitor MRTX1133. Using single-cell RNA sequencing analysis, we identified the enrichment of angiogenesis, hypoxia, and epithelial-to-mesenchymal transition (EMT) signatures in the resistant model compared to the parental PDO. Across all resistant models, VEGFA expression and VEGFR2 phosphorylation were uniformly elevated, which were driven by AKT activation and SP1 nuclear translocation. Mechanistic investigations uncovered increased PI3Kγ activity in MRTX1133-resistant models via complex formation of KRAS with p110γ and p101. This leads to an autocrine VEGFA-VEGFR2 signaling loop formation and EMT induction. Therapeutically, the disruption of VEGFA-VEGFR2 signaling restored MRTX1133 sensitivity and inhibited EMT. Furthermore, cancer-endothelial paracrine signaling amplified angiogenesis, hypoxia, and EMT signatures in cancer cells and simultaneously promoted endothelial cell proliferation, reinforcing an adaptive feedback mechanism. In a mouse model of MRTX1133-resistant tumor xenograft, a combination of anti-VEGFR2 therapy and MRTX1133 more effectively reduced tumor growth, angiogenesis, and proliferation markers than monotherapy without significant body weight change. These findings establish VEGFA-VEGFR2 signaling by PI3Kγ activation as a key driver of acquired resistance to KRAS G12D inhibition and provide a rationale for combining VEGFA-VEGFR2 inhibition with KRAS blockade in KRAS-mutant cancers.

**Highlight:**

- VEGFA-VEGFR2 signaling activation is a common feature of MRTX1133 resistance in KRAS^G12D^ cancer cells
- Nuclear translocation of SP1 by AKT activation promotes VEGFA transcription in MRTX1133-resistant models
- Interaction of p110γ-p101 with KRAS activates PI3Kγ in the resistant models
- VEGFA-VEGFR2 inhibition reverses MRTX1133 resistance *in vitro* and *in vivo*

## Introduction

Gain-of-function mutations in the *KRAS* gene constitutively activate key oncogenic signaling pathways such as PI3K-AKT, RAF, and NF-κB, promoting tumorigenesis and cancer progression (Singhal et al., 2024). Historically, KRAS mutations were considered ‘undruggable’ until the development of KRAS^G12C^-specific covalent inhibitors that have demonstrated clinical efficacy, including sotorasib and adagrasib (Ostrem et al., 2013). However, many patients have cancer harboring KRAS mutations other than G12C, necessitating the development of additional KRAS-targeted therapies. The KRAS^G12D^ mutation is highly prevalent in pancreatic (37%), gastric (13%), and colorectal (13%) cancers (Consortium et al., 2017). Recently, several inhibitors targeting KRAS^G12D^ have been developed, offering potential therapeutic opportunities for patients with these malignancies (Singhal et al., 2024).

MRTX1133 is a KRAS^G12D^-selective, non-covalent inhibitor that has demonstrated promising antitumor efficacy in preclinical studies (Hallin et al., 2022). In preclinical data, MRTX1133 combined with EGFR inhibitors exhibits synergistic antitumor effects (Mahadevan et al., 2023). In addition, the antitumor effects of MRTX1133 are partially driven by T-cells and synergize with immune checkpoint inhibitors. Investigations into MRTX1133 resistance mechanisms in preclinical models have identified EMT, PI3K-AKT pathway activation, and amplification of oncogenic drivers, including *Kras, Yap1, Myc, Cdk6, Abcb1a*, and *Abcb1b,* as contributors to acquired resistance (Dilly et al., 2024). However, therapeutic strategies to circumvent resistance to MRTX1133, especially in gastrointestinal cancer, remain insufficiently explored.

Angiogenesis plays a crucial role in tumor progression and aggressiveness by ensuring a continuous supply of oxygen and nutrients to the tumor microenvironment (Guelfi et al., 2024). Cancer cells promote endothelial cell proliferation and vascular remodeling through the secretion of pro-angiogenic factors, including vascular endothelial growth factor A (VEGFA), basic fibroblast growth factor, and platelet-derived growth factor. VEGFA binds to VEGF receptor 2 (VEGFR2, encoded in *KDR*) with high affinity, serving as a key mediator of tumor-associated angiogenesis (Simons et al., 2016). Therapeutic strategies targeting the VEGFA-VEGFR2 signaling pathway have demonstrated significant clinical efficacy. For example, ramucirumab, a monoclonal antibody against VEGFR2, has shown promising efficacy in clinical trials (Tabernero et al., 2015; Wilke et al., 2014) and is currently integrated into clinical practice.

This study aimed to identify the key resistance mechanism of MRTX1133 in KRAS^G12D^-mutant cancer cell lines and patient-derived organoids (PDOs). In this study, we found that chronic MRTX1133 exposure triggers transcriptional reprogramming: specifically, an alternative activation of PI3Kγ leading to SP1-mediated VEGFA-VEGFR2 signaling. We further reveal the molecular underpinnings of the autocrine VEGFA-VEGFR2 signaling loop in this resistance and present the overcoming strategies for MRTX1133 resistance.

## Results

### Acquired resistance to MRTX1133 is associated with the VEGFA-VEGFR2 pathway

We established MRTX1133-resistant models by treating three colorectal cancer (CRC) PDOs (SNU-4646S1-TO, SNU-6325-TO, and SNU-6330-TO), as well as two gastric cancer (GC) (AGS and SNU-601), two pancreatic cancer (PC) (AsPC-1 and PANC-1), and two CRC (SNU-C2A and SNU-C2B) cell lines for over 6 months (Supplementary Figs. S1A, S1B). Among these, parental SNU-4646S1-TO (P-PDO) and MRTX1133-resistant SNU-4646S1-TO (R-PDO) were chosen for single-cell RNA sequencing (scRNA-seq) analysis. Clustering analysis revealed two distinctly segregated groups with clusters C1, C2, and C3 predominantly composed of P-PDO, whereas clusters C6, C7, and C8 primarily consisted of R-PDO (Fig. 1A). To elucidate the mechanisms underlying acquired resistance to MRTX1133, we conducted gene set variation analysis (GSVA) using hallmark gene sets from the MSigDB (Supplementary Fig. S1C) between P-PDO-dominant clusters and R-PDO-dominant clusters. P-PDO-dominant clusters exhibited significant enrichment in cell cycle-related pathways, including E2F and MYC target gene signatures (Fig. 1B), whereas R-PDO-dominant clusters demonstrated upregulated angiogenesis, hypoxia, and epithelial-to-mesenchymal transition (EMT) gene signatures. Among genes that were increased in R-PDO-dominant clusters, nine candidates were prioritized based on the druggability (Fig. 1C). Notably, VEGFA, a key regulator of angiogenesis, hypoxia, and EMT, was selected as a potential driver of resistance.

**Figure 1.**
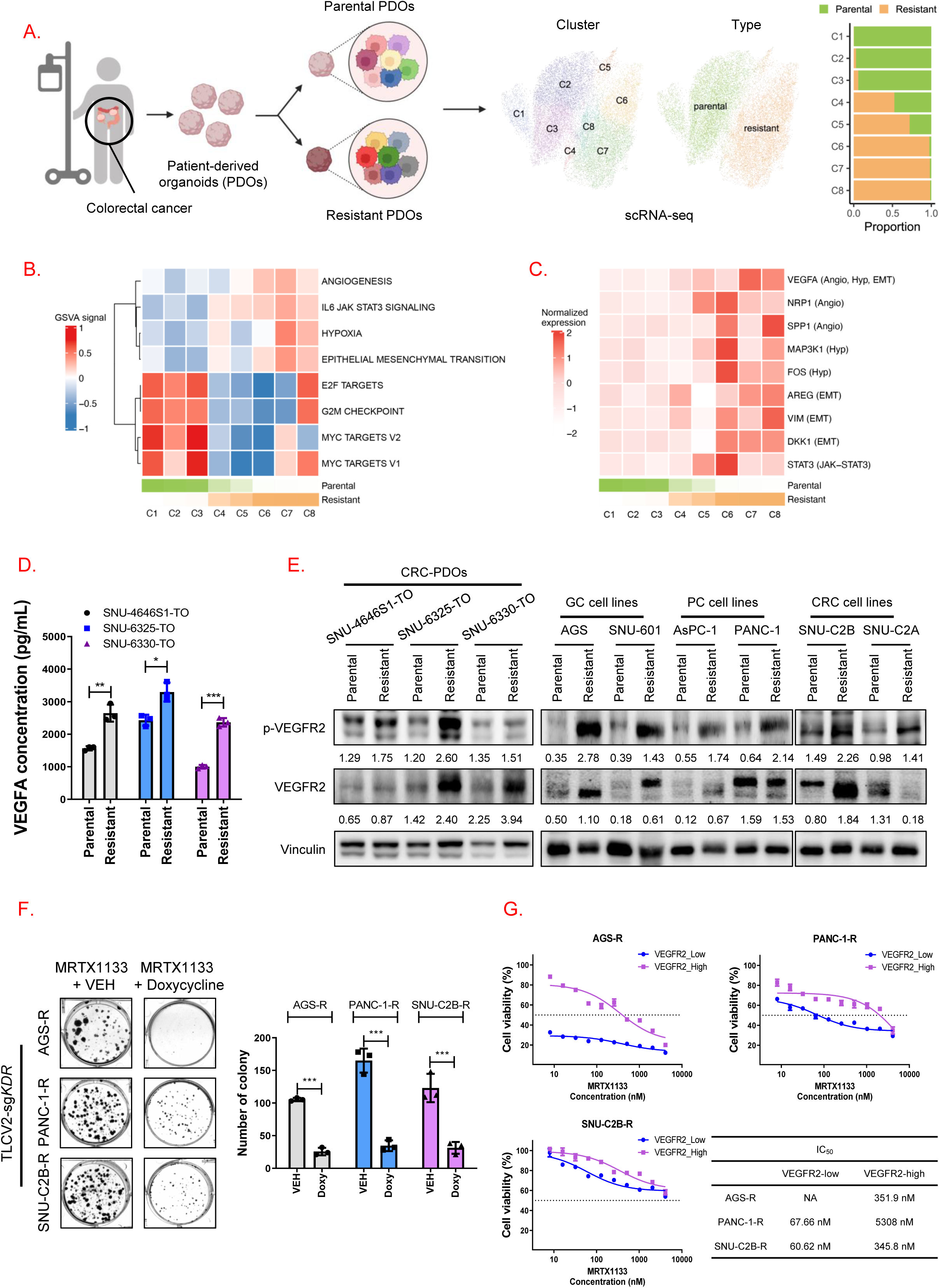
Angiogenesis driven by acquired resistance to MRTX1133. (A) Schematic diagram of generating MRTX1133-resistant cancer models and characterization of these models with a uniform manifold approximation and projection (UMAP) plot showing both parental and resistant models and the proportion of parental and resistant cells within each cluster. (B) GSVA signals of Angiogenesis, Hypoxia, Epithelial-Mesenchymal Transition, MYC Targets V2, E2F Targets, and MYC Targets V1 in the MSigDB for each cluster, with the proportions of parental and resistant cells displayed below. (C) Z-score normalized expression for nine representative genes involved in angiogenesis, hypoxia, and EMT pathways. Each gene was labeled with an abbreviation for its respective pathway: Angiogenesis (Angio), Hypoxia (Hyp), and EMT. The proportions of parental and resistant cells are shown beneath the heatmap. (D) VEGFA concentration in conditioned media (CM) derived from parental and MRTX1133-resistant patient-derived organoids (PDOs). The experiment was performed in triplicate. *P <□0.05, **P <□0.01, and ***P <□0.001. (E) Levels of p-VEGFR2, total VEGFR2, and vinculin in parental and MRTX1133-resistant models. The normalized values by vinculin were indicated. (F) Colony-forming assays were performed to assess the proliferative effect of *KDR* knockout in MRTX1133-resistant cells. TLCV2-sg*KDR-*resistant cells were treated with 1 μM of MRTX1133 and either VEH or doxycycline for 2 weeks. The colony images were captured using ChemiDoc Touch (Bio-Rad) and quantified using the ImageJ software (NIH). The Y-axis of the graph indicates the number of colonies. The experiment was performed in triplicate. ***P <□0.001. (G) The viability of VEGFR2-high and -low resistant cells sorted by FACS after MRTX1133 treatment. All experiments were repeated four times. Data represent the mean□±□SD. The IC_50_ values were calculated using GraphPad Prism Software.

To validate the upregulation of VEGFA in resistant models, we first confirmed that VEGFA protein levels were significantly higher in conditioned media (CM) derived from resistant models (R-CM) than those in their matched parental model-derived CM (P-CM) (Fig. 1D and Supplementary Fig. S1D). Treating PANC-1 cells with R-CM, compared to P-CM, enhanced VEGFR2 and p-VEGFR2 (Supplementary Fig. S1E). The p-VEGFR2 level was consistently higher, with a trend towards a higher total VEGFR2 level, in resistant models relative to their parental counterparts (Fig. 1E). To assess the functional role of VEGFR2 signaling in the MRTX1133 resistance, we inhibited VEGFR2 using ramucirumab and observed significantly suppressed proliferation of resistant cells but no apparent effect on parental cells (Supplementary Fig. S1F). To further investigate the role of VEGFR2 in the MRTX1133 resistance, we generated inducible-*KDR*-knockout (KO) MRTX1133-resistant cell lines (Supplementary Fig. S1G). Doxycycline-induced *KDR*-KO in resistant cells reduced colony formation (Supplementary Fig. S1H). When combined with MRTX1133, *KDR*-KO significantly enhanced the anti-proliferative effect compared to vehicle (VEH) control cells (Fig. 1F). Additionally, inducible-*KDR-*KO significantly reduced the VEGFA secretion (Supplementary Fig. S1I). To determine whether VEGFR2 expression levels correlated with MRTX1133 resistance, we sorted resistant cells into VEGFR2-high and VEGFR2-low populations (Supplementary Fig. S1J) and found that VEGFR2-high cells exhibited greater resistance to MRTX1133 compared to VEGFR2-low cells (Fig. 1G). Together, these data indicate that VEGFA-VEGFR2 signaling functionally contributes to MRTX1133 resistance.

### Transcriptional activation of *VEGFA* by SP1 drives acquired resistance to MRTX1133

We investigated how VEGFA is upregulated in resistant cells. Reporter assays showed significantly higher VEGFA promoter activity in resistant cells compared to parental cells (Fig. 2A). Previous studies have shown that SP1 binds to the VEGFA promoter and enhances the *VEGFA* transcription (Pore et al., 2004b). To determine whether increased VEGFA levels in resistant cells were mediated by SP1, we examined SP1 localization in nuclei. Resistant cells had markedly higher nuclear SP1 levels than parental cells (Fig. 2B, Supplementary Fig. S2A). Notably, inhibiting VEGFR2 signaling suppressed SP1 nuclear localization: ramucirumab reduced nuclear SP1 in resistant cells (Supplementary Fig. S2B), and inducible-*KDR*-KO similarly lowered SP1 expression in the nucleus (Supplementary Fig. S2C). Conversely, in parental cells, addition of recombinant human VEGFA (rhVEGFA) enhanced nuclear SP1 expression (Supplementary Fig. S2D) and abolished the responsiveness to MRTX1133(Supplementary Fig. S2E).

**Figure 2.**
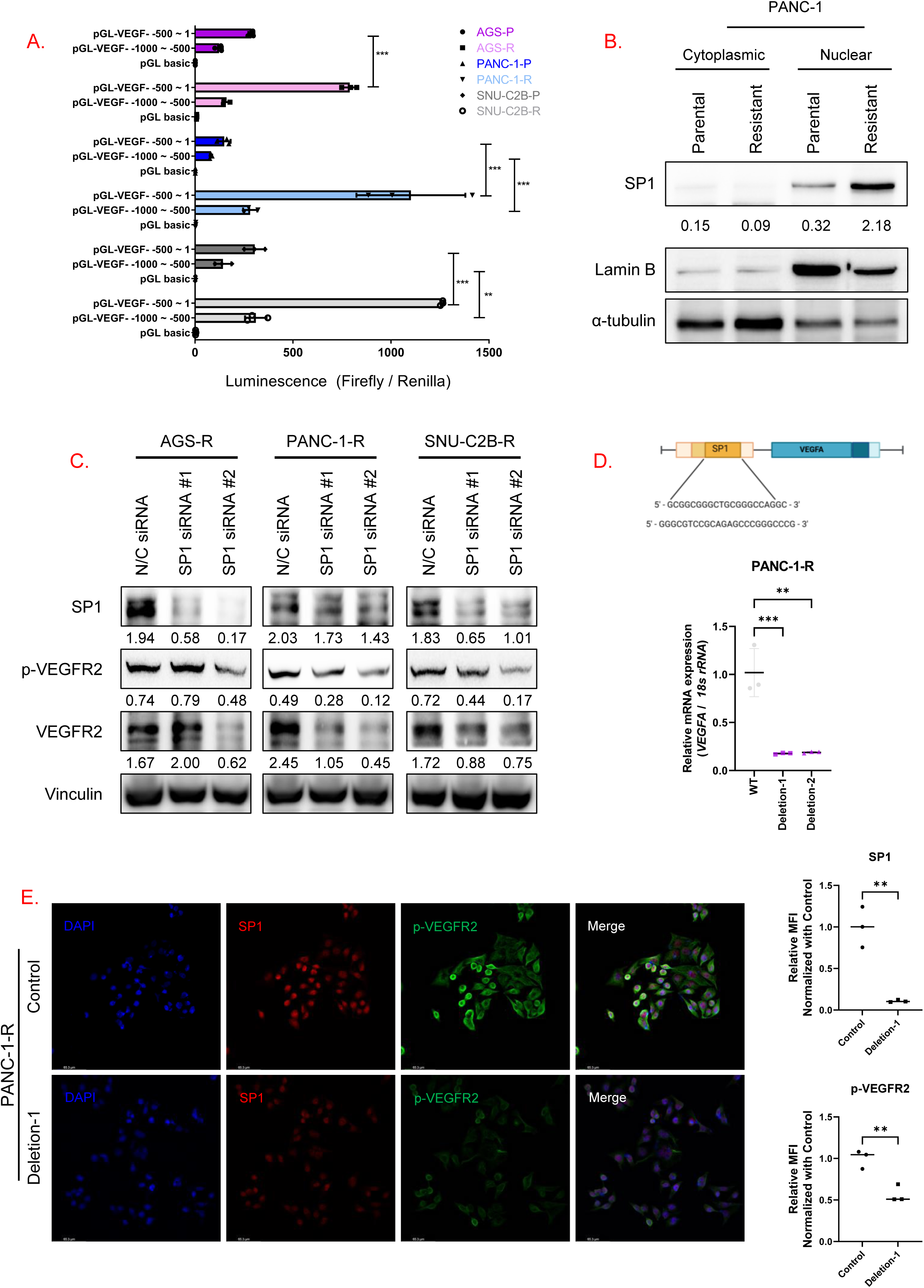
Acquired resistance to MRTX1133 driven by SP1-mediated transcriptional activation of VEGFA. (A) Relative luciferase reporter activity of two VEGFA promoter regions (1-500 bp and 500-1000 bp upstream of the transcription starting site) in parental and resistant cells. pGL basic was a negative control. The graph indicates the normalization of firefly luminescence by Renilla luminescence. All experiments were repeated three times. Data represent the mean□±□SD. **P <□0.01 and ***P <□0.001. (B) Levels of SP1, Lamin B, and α-tubulin in fractionated cytoplasmic and nuclear protein samples of PANC-1-P and PANC-1-R cells. SP1 expression was normalized by Lamin B expression. (C) Levels of SP1, p-VEGFR2, and VEGFR2 of resistant cells that were transfected with negative control (N/C) or SP1 siRNA. The values indicate the normalized band intensity by vinculin. (D) Deleting 5-bp in the predicted SP1 binding site of the *VEGFA* promoter region significantly decreased the *VEGFA* mRNA expression. *VEGFA* transcriptional levels in PANC-1-R cells transfected with wild-type and deletion vectors were plotted. The values are normalized with *18s rRNA.* All experiments were performed in triplicate. The graph indicates the mean ± SD. ***P <□0.001. (E) Representative immunofluorescence images for p-VEGFR2 (green) and SP1 (red) in PANC-1-R cells after transfection with wild-type and deletion vectors. Nuclei were stained with DAPI (blue). Scale bar: 65.3 μm.

To further investigate the role of SP1, we transfected MRTX1133-resistant cells with SP1 siRNA. Knockdown of *SP1* suppressed expression levels of p-VEGFR2 and VEGFR2 (Fig. 2C) and increased sensitivity to MRTX1133 compared to the negative control (Supplementary Fig. S2F). To verify whether SP1 directly regulated the *VEGFA* transcription, we deleted the predicted SP1 binding site in the VEGFA promoter region. SP1 binding site deletion significantly reduced *VEGFA* mRNA (Fig. 2D). Consistently, cells harboring this deletion showed lower p-VEGFR2 and nuclear SP1 expressions (Fig. 2E) compared to VEH-treated controls. These results indicate positive feedback: VEGFA-VEGFR2 signaling sustains nuclear SP1, which in turn promotes *VEGFA* transcription.

### Interaction of p110***γ***-p101 with KRAS activates PI3K***γ*** in MRTX1133-resistant cells

In line with a recent publication (Dilly et al., 2024), we found an increase in p-AKT in resistant cells compared to their parental counterparts (Supplementary Fig. S3A). Furthermore, MK2206, an AKT inhibitor, reduced nuclear SP1 levels in resistant cells (Supplementary Fig. S3B). Since RAS can activate PI3K signaling (Castellano and Downward, 2011), we speculated that enhanced interaction between KRAS and PI3K in resistant cells might trigger the autocrine VEGFA-VEGFR2-SP1 signaling loop.

To prove this, we treated both parental and resistant cells with MRTX1133 and assessed PI3K pathway activity by measuring PIP_3_ levels (Fig. 3A). In parental cells, MRTX1133 treatment significantly reduced PIP_3_ levels compared to untreated controls. In contrast, resistant cells exhibited significantly higher basal PIP_3_ levels than their parental counterparts. Even with KRAS inhibition by MRTX1133, resistant cells maintained high PI3K activity. In PANC-1-R cells with inducible-*KDR*-KO, MRTX1133 alone partially reduced VEGFR2, p-VEGFR2, and p-AKT levels (Supplementary Fig. S3C), but could not fully extinguish the signaling. Notably, the combination of MRTX1133 and doxycycline effectively abrogated the vicious loop of VEGFA-VEGFR2 signaling.

**Figure 3.**
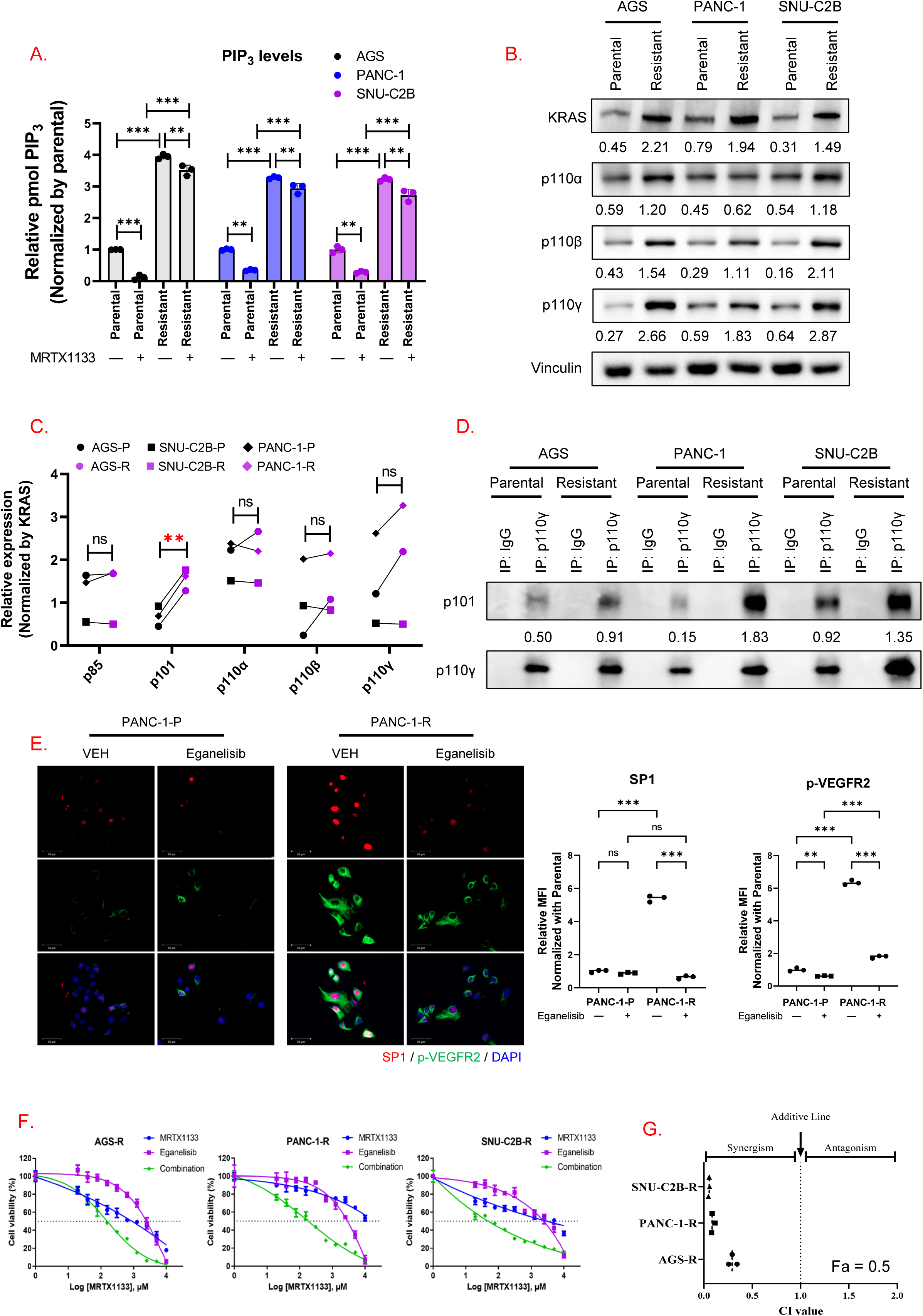
PI3Kγ activation by increased interaction of p110γ-p101 with KRAS in resistant cells. (A) Levels of PtdIns (3,4,5) P_3_ of parental and resistant cells following treatment with VEH or 1 μM MRTX1133. All experiments were repeated three times. Data represent the mean□±□SD. ***P <□0.001. (B) Levels of KRAS, p110α, p110β, p110γ, and vinculin in parental and resistant cell lines. The values indicate the normalized band intensity by vinculin. (C) The KRAS protein in parental and resistant cells is immunoprecipitated, and the p85, p101, p110α, p110β, and p110γ protein levels are analyzed. The values were normalized by KRAS. The graph indicated the normalized value of AGS, SNU-C2B, and PANC-1 parental and resistant cells. **P <□0.01. (D) p110γ proteins in parental and resistant cells were immunoprecipitated, and p101 protein levels were analyzed. The values of p101 were normalized by p110γ. (E) Representative immunofluorescence images for p-VEGFR2 (green) and SP1 (red) in PANC-1 parental and resistant cells after treatment with 1 μM of eganelisib. Nuclei were stained with DAPI (blue). Scale bar: 50 μm. Relative MFI value of SP1 and p-VEGFR2. **P <□0.01 and ***P < 0.001. (F) The viability of resistant cells after treatment with MRTX1133 and/or eganelisib. The experiment was repeated four times. Error bars indicate the mean□±□SD. (G) Combination index values were calculated after the combination treatment of 10 μM MRTX1133 and 10 μM eganelisib in resistant cells. All experiments were performed in triplicate. Data represent the mean□±□standard error of the mean (SEM).

Then, we aimed to further examine the expression levels of KRAS and PI3K catalytic subunits, including class IA (p110α and p110β) and class IB (p110γ), in whole-cell lysates, and found that the expression levels of KRAS, p110β, and p110γ were elevated in resistant cells compared with parental cells (Fig. 3B). Next, we examined the interaction between KRAS and PI3K catalytic subunits including p110α, p110β, and p110γ, and regulatory subunits, p85 and p101 (Sun and Meng, 2020). Co-immunoprecipitation studies uncovered that KRAS binds much more to p101 in resistant cells than in parental cells, whereas binding to other PI3K components (p85, p110α/β/γ) remained unchanged (Fig. 3C and Supplementary Fig. S3D). In previous studies (Rathinaswamy et al., 2021; Rathinaswamy et al., 2023), the p110γ-mediated p101 and RAS complex substantially boosts PI3Kγ activity compared to the monomeric or dimeric forms. Based on this, we evaluated the interaction between p110γ and p101 in both parental and resistant cells. Consistently, we observed enhanced p110γ-p101 interaction in resistant cells (Fig. 3D).

By targeting PI3Kγ with eganelisib, a selective p110γ inhibitor, we sought to break the autocrine loop. Eganelisib treatment of resistant cells markedly reduced AKT activation and VEGFA secretion (Supplementary Figs. S3E and S3F), also lowering nuclear SP1 and p-VEGFR2 levels (Fig. 3E). Functionally, combining eganelisib with MRTX1133 synergistically suppressed the viability of resistant cells (Figs. 3F, 3G, and Supplementary Fig. S3G), confirming that PI3Kγ activity is critical for the survival of MRTX1133-resistant cells.

Resistant cells exhibited higher transcriptional levels of *KRAS* and *PIK3CG* than those of parental cells (Supplementary Fig. S3H). Next, to determine whether the increased expression of KRAS and p110γ in resistant cells was attributed to altered protein stability, we assessed their expression following treatment with cycloheximide (CHX) and MG132, inhibitors of protein synthesis and proteasomal degradation, respectively (Supplementary Fig. S3I). After CHX treatment, KRAS and p110γ levels in resistant cells initially increased but then declined, indicating limited stability when new synthesis is halted. In parental cells, both proteins remained low and largely unchanged. With MG132, p110γ robustly accumulated in resistant cells, consistent with proteasome-mediated degradation, whereas KRAS slightly decreased despite proteasome inhibition. In contrast, parental cells showed clear KRAS accumulation upon proteasome blockade. Taken together, elevated KRAS and p110γ levels in resistant cells are driven primarily by transcriptional upregulation rather than reduced proteasomal degradation. Moreover, p110γ in resistant cells behaves as a proteasome-sensitive, high-turnover protein whose higher steady-state level reflects increased transcription counterbalanced by active degradation, whereas KRAS abundance is largely transcription-driven with little evidence of proteasomal stabilization.

### Autocrine VEGFA-VEGFR2 loop inhibition reverses EMT in MRTX1133-resistant cells

Based on the enrichment of EMT-associated gene sets in the scRNA-seq analysis (Fig. 1B), we hypothesized that the PI3Kγ-driven VEGFA-VEGFR2-SP1 loop contributes to an EMT-like phenotype in resistant cells. To assess the role of PI3Kγ activity in promoting the mesenchymal state, we performed migration assays following treatment with eganelisib, a selective p110γ inhibitor. In resistant cells, eganelisib treatment significantly reduced cell migration compared to VEH (Fig. 4A). Additionally, eganelisib treatment suppressed the expression of SP1 and mesenchymal markers, including ZEB1 and vimentin, in resistant cells (Fig. 4B). To further validate the role of SP1 on EMT, we knocked down *SP1* using siRNA and found that *SP1* knockdown significantly reduced cell migration (Fig. 4C) and mesenchymal markers expression (Fig. 4D) in resistant cells. As shown previously (Fig. 2C and Supplementary Fig. S3F), the autocrine VEGFA-VEGFR2 signaling loop was regulated by both SP1 and PI3Kγ. To determine whether the EMT phenotype in resistant cells is attributed to VEGFR2 activation, we conducted migration assays following VEGFR2 inhibition. In resistant cells, doxycycline-induced *KDR* KO significantly reduced the number of migrated cells compared to VEH treatment (Fig. 4E). Similarly, ramucirumab treatment inhibited migration in resistant cells, in contrast to parental cells (Fig. 4F). Collectively, the autocrine VEGFA-VEGFR2 signaling loop drives not only survival under KRAS inhibition but also EMT and motility.

**Figure 4.**
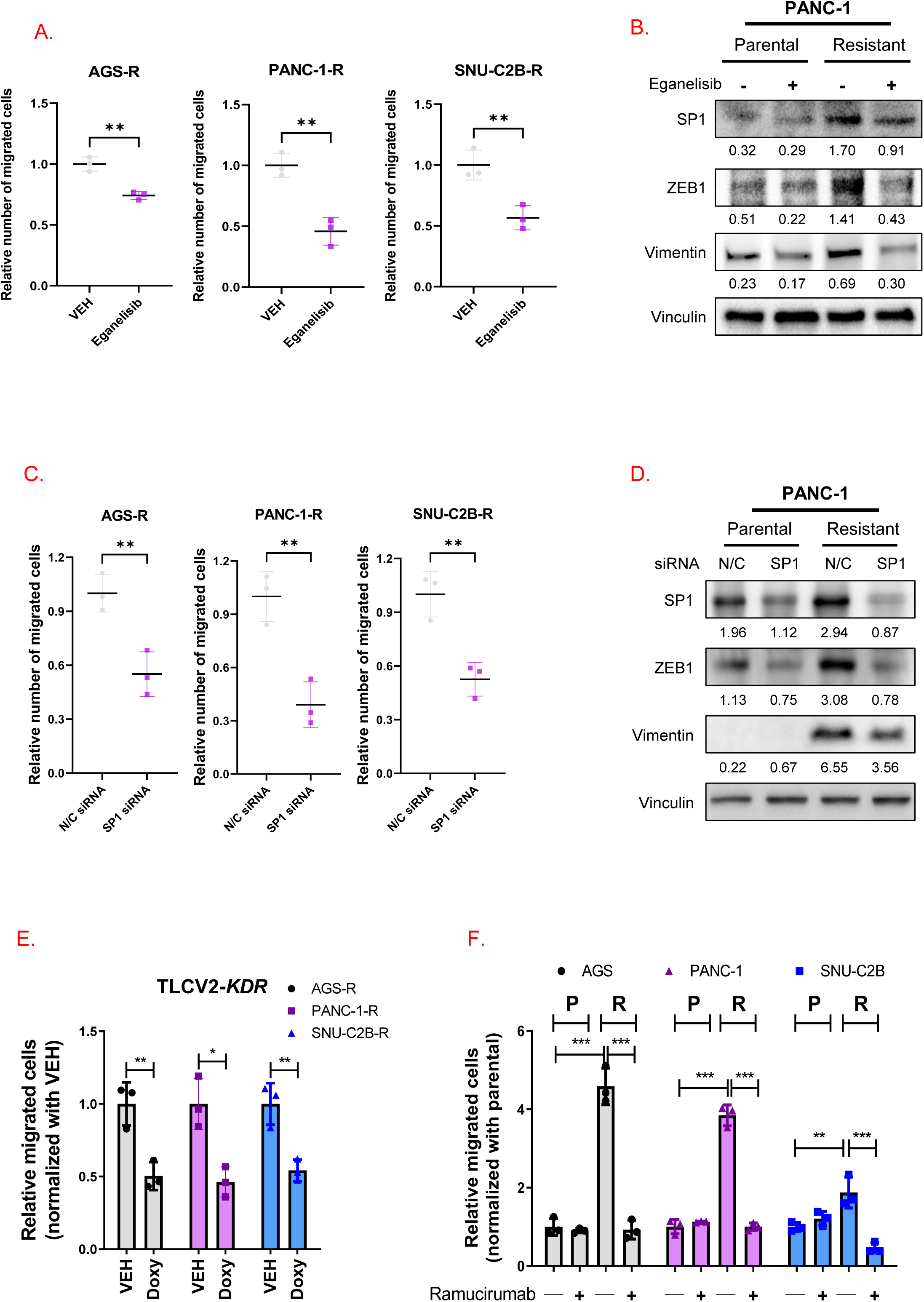
Effect of VEGFA-VEGFR2-PI3Kγ-SP1 signaling loop on EMT in MRTX1133-resistant cells. (A) The number of migrated MRTX1133-resistant cells treated with VEH and eganelisib. The graph indicates the number of migrated cells relative to those treated with VEH. The experiment was independently performed three times. **P <□0.01. (B) Levels of ZEB1, vimentin, and vinculin in PANC-1-P and PANC-1-R cells treated with 1 μM eganelisib. The values represent the normalized band intensities by vinculin. (C) The number of migrated MRTX1133-resistant cells transfected with siRNA for *SP1* relative to those transfected with the negative control siRNA. The experiment was independently performed three times. **P <□0.01. (D) Levels of SP1, ZEB1, vimentin, and vinculin in PANC-1-R cells after transfection with siRNA for negative control or *SP1.* The value indicates the normalized values by vinculin intensity. (E) The number of migrated *KDR*-KO resistant cells by TLCV2-sg*KDR* after treatment with doxycycline relative to those treated with VEH. The experiment was independently performed three times. *P <□0.05 and **P <□0.01. (F) The number of migrated parental and resistant cells after treatment with 50 μg/mL ramucirumab. The experiment was independently performed three times. **P <□0.01 and ***P <□0.001.

### Cancer-endothelial paracrine interaction amplifies tumor microenvironment dynamics

Given the differential tube formation of HUVECs in response to R-CM versus P-CM (Supplementary Fig. S4A), we investigated the paracrine interaction between HUVECs and resistant cells. When co-cultured with HUVECs, PANC-1-R cells more significantly co-localized with HUVECs than PANC-1-P cells (Figs. 5A, 5B, and Supplementary Fig. S4B). Additionally, PANC-1-R cell proliferation was significantly increased when cultured in CM from HUVECs (HUVEC-CM) compared to complete media (Supplementary Fig. S4C). To uncover the molecular mechanisms driving tumor-endothelial interactions, we further characterized the interactions between PDOs and HUVECs using indirect co-culture of PDOs (parental or MRTX1133-resistant SNU-4646S1-TO) and HUVECs. We performed bulk RNA sequencing (RNA-seq) on four PDO groups and three HUVEC groups. The four PDO groups were (1) P-PDO without HUVEC co-culture (P-PDO^-HUVEC^), (2) R-PDO without HUVEC co-culture (R-PDO^-HUVEC^), (3) P-PDO with indirect HUVEC co-culture (P-PDO^+HUVEC^), and (4) R-PDO with indirect HUVEC co-culture (R-PDO^+HUVEC^). The three HUVEC groups were (1) HUVECs without PDO co-culture (HUVEC^-PDO^), (2) HUVECs with indirect P-PDO co-culture (HUVEC^+P-PDO^), and (3) HUVECs with indirect R-PDO co-culture (HUVEC^+R-PDO^).

**Figure 5.**
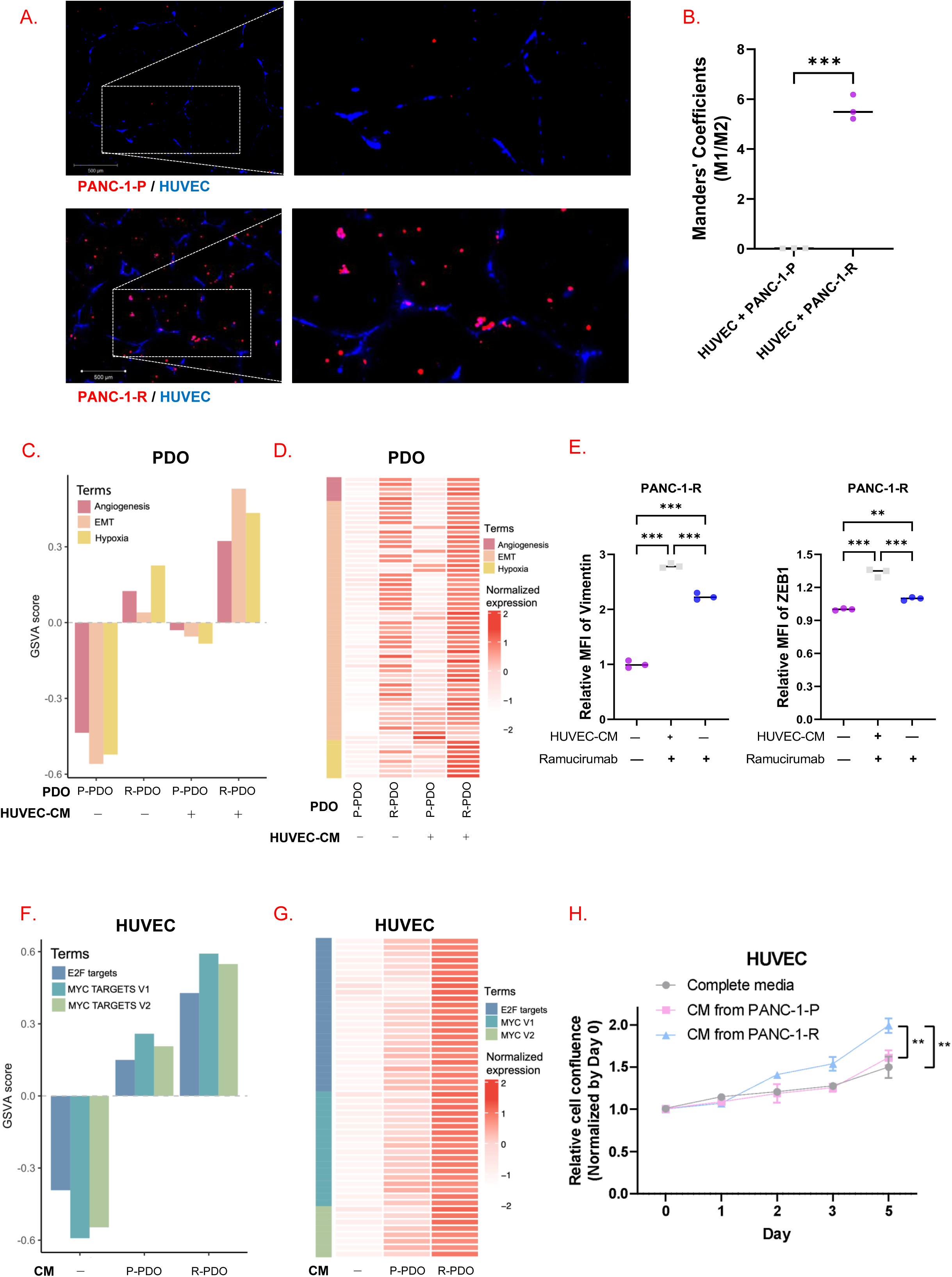
Interaction between MRTX1133-resistant cells and HUVECs. (A) Representative fluorescence images of HUVECs (blue) and cancer cells (red) seeded in Matrigel-precoated plates and incubated for 18 h. Scale bar: 200 μm. (B) Quantitative co-localization analysis of HUVEC and PANC-1-P or PANC-1-R. Mander’s coefficients M1 (fraction of HUVEC signal overlapping PANC-1-P or PANC-1-R) and M2 (fraction of PANC-1-P or PANC-1-R signal overlapping HUVEC) were calculated using the Coloc2 plugin in Fiji (ImageJ, NIH). ***P <□0.001. (C) GSVA scores for the Angiogenesis, EMT, and Hypoxia pathways in each PDO. (D) Expression of individual genes in the Angiogenesis, EMT, and Hypoxia pathways. (E) Relative MFI value of vimentin and ZEB1 in PANC-1-R cells treated with complete media, CM from HUVEC, and CM from HUVEC combined with ramucirumab. **P <□0.01 and ***P <□0.001. (F) GSVA scores for E2F TARGETS, MYC TARGETS V1, and MYC TARGETS V2 pathways in each HUVEC. (G) Gene expression of individual genes in E2F TARGETS, MYC TARGETS V1, and MYC TARGETS V2 pathways. (H) The number of HUVECs treated with complete media, P-CM, and R-CM for 5 days. Data are presented as the mean ± SD. The experiment was independently carried out five times. **P <□0.01.

Bulk RNA-seq of the PDOs revealed that, compared to other PDOs (P-PDO^+HUVEC^, P-PDO^-HUVEC^, and R-PDO^-HUVEC^), R-PDO^+HUVEC^ exhibited the highest enrichment of EMT, angiogenesis, and hypoxia signatures (Fig. 5C), along with increased expression of individual genes (Fig. 5D and Supplementary Fig. S4D). Functionally, compared to cells without HUVEC-CM co-culture, PANC-1-R cells under HUVEC-CM displayed elevated mean fluorescence intensity (MFI) of mesenchymal markers, including ZEB1 and vimentin (Fig. 5E). Importantly, this HUVEC-CM-induced EMT phenotype was blocked by ramucirumab. These results suggested that enhanced VEGFR2 signaling by indirect HUVEC co-culture contributed to the mesenchymal phenotype of resistant cells.

A similar analysis of HUVEC transcriptional profiles revealed that, compared to other HUVECs (HUVEC^-PDO^ and HUVEC^+P-PDO^), HUVEC^+R-PDO^ upregulated genes in E2F and MYC target gene sets, key regulators of cell proliferation (Figs. 5F, 5G, and Supplementary Fig. S4E). Functionally, HUVEC proliferation was significantly enhanced when cultured under R-CM, compared to complete media or P-CM (Fig. 5H). These findings indicate that paracrine signaling between MRTX1133-resistant cells and endothelial cells promotes EMT, angiogenesis, and hypoxia signatures in cancer cells, while simultaneously enhancing endothelial proliferation through transcriptional reprogramming. This bidirectional crosstalk might reinforce tumor progression and drug resistance.

### VEGFR2 blockade overcomes acquired MRTX1133 resistance *in vivo*

To assess the therapeutic potential of VEGFR2 inhibition in MRTX1133-resistant tumors, PANC-1-P and PANC-1-R xenograft mouse models were randomized to receive VEH, DC101 (an anti-mouse VEGFR2 antibody), MRTX1133, or DC101 combined with MRTX1133. In the PANC-1-P model, both MRTX1133 monotherapy and DC101 monotherapy significantly reduced tumor growth (Supplementary Fig. S5A). However, the tumor growth inhibition (TGI) of DC101 monotherapy was lower than that of MRTX1133 alone (Supplementary Fig. S5B), and combination therapy did not provide a significant benefit over either monotherapy.

In contrast, in the PANC-1-R model, DC101 monotherapy initially induced significant suppression of tumor growth; however, tumor regrowth was observed after 15 days of treatment (Fig. 6A). The antiproliferative effect of MRTX1133 was less potent than that of DC101. Notably, the combination of MRTX1133 and DC101 exerted a marked tumor growth inhibition compared to monotherapy, achieving 85% TGI by day 15 (Fig. 6B). The mouse body weight remained stable across all treatment groups (Supplementary Fig. S5C). Moreover, the expression levels of p-VEGFR2, p-AKT, and CD31, an endothelial cell marker, were stronger in VEH- and MRTX1133-treated tumors than in DC101- or combination-treated tumors (Figs. 6C and 6D). Furthermore, the combination therapy decreased Ki67, a proliferation marker, and endomucin (ENCN), a capillary endothelial marker, more effectively than either DC101 or MRTX1133 monotherapy (Fig. 6E).

**Figure 6.**
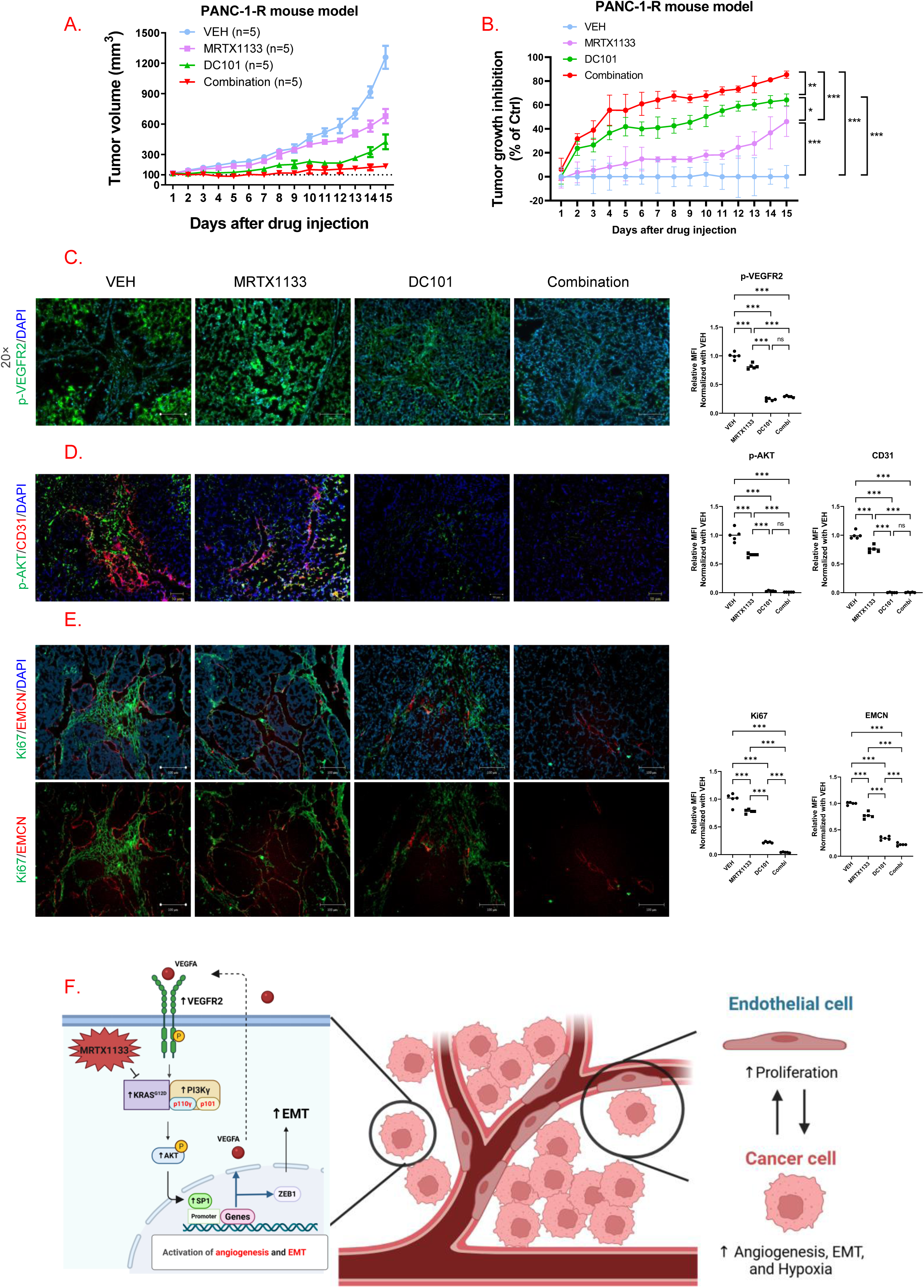
VEGFR2 blockade to overcome acquired resistance to MRTX1133 *in vivo*. (A) Tumor volumes of PANC-1-R cell-injected mice treated as indicated (n = 5 in each group). They were recorded for 15 days. (B) In the resistant model receiving MRTX1133 and/or DC101 for 15 days, the tumor growth inhibition (TGI) ratio was calculated relative to the VEH value. Data represent the mean□±□SD (n = 5). *P <□0.05, **P <□0.01, and ***P <□0.001. (C) Representative immunofluorescence images for p-VEGFR2 (green) in tumor slides from the resistant models receiving indicated treatments. Nuclei were stained with DAPI (blue). Scale bar: 100 μm. (D) Representative immunofluorescence images for p-AKT (green) and CD31 (red) in tumor slides from the resistant models receiving indicated treatments. Nuclei were stained with DAPI (blue). Scale bar: 50 μm. (E) Representative immunofluorescence images for Ki67 (green) and EMCN (red) in tumor slides from the resistant models receiving indicated treatments. Nuclei were stained with DAPI (blue). Scale bar: 100 μm. (F) A schematic summarizing the underlying mechanism of MRTX1133 acquired resistance in our models.

Collectively, our findings indicate that acquired resistance to MRTX1133 is driven by increased PI3Kγ activity, which initiates a self-sustaining autocrine VEGFA-VEGFR2 signaling loop via AKT activation and SP1 nuclear translocation (Fig. 6F). Cancer-endothelial paracrine signaling promotes angiogenesis, hypoxia, and EMT signatures in cancer cells and proliferation of vascular endothelial cells simultaneously. Inhibition of the VEGFA-VEGFR2 signaling loop reverses MRTX1133 resistance *in vitro* and *in vivo*.

## Discussion

MRTX1133 has shown promising preclinical anti-tumor effects in GC, CRC, and PDAC harboring the KRAS^G12D^ mutation (Hallin et al., 2022). However, like other anti-cancer drugs, acquired resistance is inevitable. Two principal mechanisms underlie resistance to KRAS inhibitors. Genetic resistance mechanism arises from secondary alterations within the RAS signaling cascade as observed in patient-derived samples (Awad et al., 2021; Dilly et al., 2024). The other mechanism is transcriptional reprogramming that promotes cellular plasticity, including activation of the PI3K-AKT-mTOR pathway and EMT—a phenomenon reproducibly observed in KRAS-mutant preclinical models (Dilly et al., 2024). In this study, we uncovered a common, but previously unrecognized, mechanism of acquired resistance to MRTX1133 across gastric, pancreatic, and colorectal cancer cells: alternative activation of PI3Kγ driving SP1-mediated VEGFA-VEGFR2 signaling loop.

A recent study by Kemp and colleagues reported that MRTX1133-treated *Kras*^G12D^ and *Trp53*^R172H^ (KPC) mouse models exhibited increased vascular development within the tumor mass after 7 days of treatment (Kemp et al., 2023). However, the underlying mechanisms of this phenomenon remained undiscovered. More recently, Dilly and colleagues identified angiogenesis as a hallmark feature in the mesenchymal cell population of MRTX1133-resistant models. Consistent with these findings, our scRNA-seq analysis demonstrated enrichment of angiogenesis gene sets, including the *VEGFA* gene, in MRTX1133-resistant models. We validated consistent elevation of VEGFA expression and p-VEGFR2 across six MRTX1133-resistant cell lines (two each from gastric, colorectal, and pancreatic cancer) and in three resistant colorectal PDOs, indicating that autocrine VEGFA-VEGFR2 signaling is a common feature of resistance in MRTX1133-resistant models (Fig. 1F). Emerging evidence suggests that PI3K/AKT signaling enhances SP1 nuclear translocation to bind to the proximal promoter of the *VEGF* gene (Pore et al., 2004a; Su et al., 2017). Our study revealed that VEGFR2 inhibition led to p-AKT inactivation and decreased intranuclear SP1 expression. In addition, SP1 silencing restored MRTX1133 sensitivity in resistant cells. Notably, the dual inhibition of VEGFR2 signaling and KRAS successfully overcame resistance in both *in vitro* and *in vivo* models. These findings provide a rationale for further clinical studies of combination strategies involving VEGFR2 inhibitors and KRAS inhibitors in patients experiencing resistance after KRAS inhibitor treatment.

We then identified that enhanced PI3Kγ activity triggers the acquired resistance. Direct interaction between RAS protein and p110γ through the RAS-binding domain (RBD) contributes to potent PI3K-class IB activation (Rathinaswamy et al., 2023). Although KRAS expression was elevated in whole lysates of resistant cells, its direct binding to p110 catalytic subunits (p110α, p110β, and p110γ) did not differ significantly from that observed in parental cells. By contrast, we observed a markedly stronger interaction between KRAS and p101, a regulatory subunit of p110γ, whereas the binding of p85, a canonical regulatory subunit of class IA PI3Ks, remained unchanged. Previous studies have shown that increased expression of p110γ enhances its association with p101, thereby potentiating PI3Kγ activity (Perino et al., 2011). Moreover, the Ras-p110γ-p101 tri-complex has been reported to promote PI3K signaling more effectively than mono- or dimeric assemblies (Rathinaswamy et al., 2023; Vadas et al., 2013b). In this context, p110γ may function as a molecular scaffold that bridges KRAS and p101, enabling tri-complex formation and thereby amplifying PI3Kγ signaling activation, which in turn drives the autocrine VEGFA-VEGFR2 signaling loop in MRTX1133-resistant models. Furthermore, we found that the increase of KRAS protein expression in resistant cells was attributed to increased mRNA transcription, rather than decreased protein degradation. Further studies are warranted to identify the molecular mechanism by which continuous exposure to MRTX1133 drives the increased KRAS transcription and promotes KRAS-PI3K interactions.

Eganelisib, a p110γ inhibitor, has demonstrated promising results in combination with immune checkpoint inhibitors in patients with triple-negative breast cancer (O’Connell et al., 2024). Although our *in vitro* data demonstrated that p110γ inhibition restored MRTX1133 sensitivity in resistant cells, clinical evidence of eganelisib treatment remains limited in other KRAS-driven malignancies, including pancreatic, colorectal, and gastric cancers. In contrast, ramucirumab has established clinical indications and combination protocols in patients with colorectal and gastric cancers (Tabernero et al., 2015; Wilke et al., 2014). Based on this rationale, in combination with MRTX1133, DC101, an anti-VEGFR2 antibody, was chosen for *in vivo* experiments. This approach could be readily tested in patients with gastrointestinal cancers, given that anti-VEGF/VEGFR therapies are already part of the clinical armamentarium for these diseases.

Our study further demonstrated that paracrine interactions between MRTX1133-resistant cancer cells and endothelial cells amplify angiogenesis and EMT in cancer cells. HUVEC-CM enhanced EMT in PANC-1-R cells (Figs. 5C-E), consistent with prior studies showing that HUVEC-CM upregulates mesenchymal markers, including MMP-2, MMP-9, and vimentin (Ou et al., 2019). Conversely, VEGFR2 blockade suppressed EMT in resistant cells treated with HUVEC-CM, confirming that tumor-endothelial interactions reinforce the mesenchymal state. Additionally, HUVECs treated with R-CM enhanced MYC and E2F gene signatures and endothelial proliferation. These findings indicate that resistant cancer cells and endothelial cells engage in reciprocal transcriptional reprogramming, promoting each other’s growth. Therefore, blocking this tumor-endothelial crosstalk by VEGFR2 inhibitors could be a promising strategy to overcome MRTX1133 resistance.

Many KRAS inhibitors, including MRTX1133, have been developed recently, some currently undergoing clinical trials (Cox et al., 2025). Further research is needed to determine whether this resistance mechanism is replicated in the other KRAS inhibitors. This mechanism mirrors resistance patterns seen in the *BRAF*^V600E^ inhibitor class, where both vemurafenib and dabrafenib consistently induce compensatory MEK upregulation and downstream MAPK pathway reactivation (Caeser et al., 2019; Long et al., 2014), which were overcome by the MEK inhibitor combination. This example underscores that parallel or compensatory pathways can sustain tumor growth when an oncoprotein is pharmacologically silenced and highlights how common resistance mechanisms within an inhibitor class could be effectively druggable by a combination strategy.

In summary, our study reveals a novel resistance mechanism to MRTX1133, in which alternative activation of PI3Kγ induces SP1 upregulation, driving a self-reinforcing loop of VEGFA-VEGFR2 signaling and EMT. Importantly, the concurrent inhibition of VEGFA-VEGFR2 signaling and KRAS effectively overcame resistance in preclinical models. These findings provide a rationale for clinically evaluating the combined VEGFA-VEGFR2 and KRAS inhibition in patients resistant to KRAS-targeted therapy. Future studies should focus on validating these mechanisms in clinical trials and identifying biomarkers that predict which patients benefit more from adding VEGFR2 or PI3Kγ inhibitors to KRAS inhibitor regimens.

## Methods

### Patient-derived cancer models and reagents

A total of nine patient-derived *KRAS*^G12D^-mutant cancer models were used in this study: six cell lines including gastric cancer (GC) (AGS and SNU-601), pancreatic cancer (PC) (PANC-1 and AsPC-1), and colorectal cancer (CRC) (SNU-C2A and SNU-C2B), and three CRC PDOs (SNU-4646S1-TO (Kim et al., 2022), SNU-6325-TO, and SNU-6330-TO). All patient-derived models were purchased from the Korean Cell Line Bank (Seoul, Republic of Korea). AGS, SNU-601, AsPC-1, SNU-C2A, and SNU-C2B cell lines were cultured in RPMI-1640 medium (WELGENE, Gyeongsan, Republic of Korea) supplemented with 10% fetal bovine serum (FBS) (Invitrogen, Carlsbad, CA, USA) and 1% penicillin-streptomycin (Corning, NY, USA). PANC-1 cell line was cultured in DMEM containing 10% FBS (Invitrogen) and 1% penicillin-streptomycin (Corning). MRTX1133 was purchased from MedChemExpress (Monmouth Junction, NJ, USA). Cycloheximide and eganelisib were purchased from Selleckchem (Houston, TX, USA). Recombinant human VEGFA was purchased from R&D Systems (Minneapolis, MN, USA). DC101, an anti-mouse VEGFR2 antibody, was purchased from Bio X Cell (Lebanon, NH, USA).

### Development of MRTX1133-resistant models

Patient-derived, MRTX1133-resistant, *KRAS*^G12D^-mutated cancer models were generated by treating AGS, SNU-601, PANC-1, AsPC-1, SNU-C2A, SNU-C2B, SNU-4646S1-TO, SNU-6325-TO, and SNU-6330-TO with MRTX1133. The half-maximal inhibitory concentration (IC_50_) of MRTX1133 for each model was determined using the CellTiter-Glo assay (Promega, Madison, WI, USA). Following MRTX1133 treatment, the surviving cells were cultured in a drug-free medium for 3 days. The dosages were gradually escalated from IC_6.25_ to IC_50_ over time. After 6 months, the IC_50_ values for MRTX1133-resistant cell lines exceeded 1 μM, confirming the development of acquired resistance.

### Processing of PDO and public scRNA-seq data

Single-cell RNA sequencing (scRNA-seq) libraries were generated for parental (SNU-4646S1-TO-P) and MRTX1133-resistant (SNU-4646S1-TO-R) patient-derived organoids (PDOs) using the Chromium Next GEM Single Cell 3’ Reagent Kits v3.1 (10x Genomics, Pleasanton, CA, USA). Sequencing was performed on the Illumina HiSeq X system (San Diego, CA, USA), and .fastq files were preprocessed using Cell Ranger v7.0.1 (10x Genomics). Reads were aligned to the GRCh38 reference genome (GRCh38-2020-A), including intronic regions. Subsequent analyses were conducted using Scanpy v1.10.1 (Wolf et al., 2018). For quality control, cells were filtered based on the following criteria: (1) number of detected genes < 200, (2) total unique molecular identifiers (UMIs) < 10,000, and (3) mitochondrial content > 20%. Doublets were removed using Scrublet (Wolock et al., 2019) within Scanpy. Highly variable genes (n = 2,000) were identified using the Pearson residual method. For clustering analysis, expression values were normalized using Pearson residuals. Log-transformed counts per million (logCPM) were used for expression visualization and Gene Set Variation Analysis (GSVA). Clustering was performed using the top 40 principal components, followed by neighborhood graph construction with default parameters and cluster identification via the Leiden algorithm at a resolution of 0.4.

### Processing of bulk RNA-seq in the cancer and endothelial cell co-culture system

Organoids were enzymatically and mechanically dissociated into single cells by incubating them in TrypLE Express solution (Gibco, Thermo Fisher Scientific, Waltham, MA, USA) for 5–10 minutes with gentle pipetting. The dissociated cells were then neutralized with fetal calf serum (FCS) and complete medium before centrifugation at 1,500 rpm for 3 minutes. The cell pellet was washed once with FCS and complete medium, resuspended in basement membrane extract (BME), and seeded as 50–100 μL domes on the insert portion of a Co-Culture Dish (SPL Life Sciences, Pocheon-si, Republic of Korea; Cat#209260). After BME solidification, Human Intestinal Stem Cell (HISC) medium was added to overlay the BME domes, and organoids were incubated at 37°C in a 5% COL incubator. Organoids were stabilized in HISC medium for approximately 4 days, allowing them to reach 70% confluency.

At 70–90% confluency, Human Large Vessel Endothelial Cells (HUVECs) were detached using trypsin, centrifuged, and resuspended in HUVEC culture medium. A total of 3 × 10L HUVECs were seeded on the bottom of the Co-Culture Dish in 1 mL of Human Large Vessel Endothelial Cell Basal Medium (Gibco) after removal of HISC medium. The co-culture was incubated at 37°C in a 5% COL incubator using HUVEC medium.

After 4 days of co-culture, organoids and HUVECs were dissociated into single cells using TrypLE Express (for organoids) and trypsin (for HUVECs), then washed three times with PBS. The resulting cell pellets were stored in TRIzol (Invitrogen, Carlsbad, CA, USA) for RNA sequencing.

For the control group, organoids and HUVECs were seeded in the insert and bottom portions of the Co-Culture Dish, respectively, but maintained in separate culture dishes under identical conditions. Cells were harvested as pellets following the same dissociation protocol as the experimental group.

### Processing of bulk RNA-seq

Bulk RNA sequencing (RNA-seq) was performed on the NovaSeq 6000 system (Illumina) using RNA extracted from the PDOs and HUVECs co-culture system. Raw sequencing data were converted from .bcl to .fastq format, and low-quality reads were filtered using Fastp version 0.23.4 (Chen et al., 2018). Filtered reads were aligned to the GRCh38 reference genome using the STAR pipeline v2.7.11b (Dobin et al., 2013) with “Basic” two-pass mode and Gencode V44 annotations. Duplicate reads were marked using Picard version 3.1.1, and gene expression was quantified using featureCounts version 2.0.6. Quality control metrics were assessed using Qualimap version 2.3 [26428292]. Gene expression levels were calculated as transcripts per million (TPM) using the following formulas:

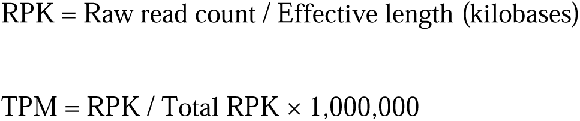

### GSVA analysis for scRNA-seq and bulk RNA-seq

GSVA was performed using the gseapy Python library (v1.1.3) (Fang et al., 2023) and the human Molecular Signatures Database (MSigDB, 2023.2.Hs). LogCPM values from scRNA-seq and TPM values from bulk RNA-seq were used as input data for the analysis.

### Cell sorting according to VEGFR2 expression

MRTX1133-resistant cells were stained with an anti-VEGFR2 antibody (Santa Cruz Biotechnology, CA, USA), followed by an incubation with an Alexa Fluor 647-conjugated secondary antibody (Invitrogen). Stained cells were then sorted into VEGFR2-high and VEGFR2-low populations using BD FACS Aria III (BD Bioscience, Franklin Lakes, NJ, USA). The sorted cells were cultured in complete media and subsequently used for further experiments.

### Calculation of IC_50_ and proliferation

Cell viability was assessed following MRTX1133 treatment using the CellTiter-Glo reagent (Promega) per the manufacturer’s instructions. The IC_50_ value was determined using GraphPad Prism (GraphPad, Boston, MA, USA). For proliferation assays, cells were seeded in 24-well plates and monitored for 5 days using the NYONE imaging system (SYNENTEC, Elmshorn, Germany). Proliferation values were normalized to day 0 for relative comparison.

### Generation of genetically-engineered cells

Inducible KDR knockout (KO) cell lines were generated using the CRISPR/Cas9 system. Single-guide RNA (sgRNA) sequences targeting *KDR* (forward: CACCGTTACCCGCTTAACGGTCCGTAGG, reverse: AAACCCTACGGACCGTTAAGCGGGTAAC) were designed using CHOPCHOP (https://chopchop.cbu.uib.no/) and cloned into the TLCV2 vector (Addgene, #87360) following *BSMBI* digestion.

For lentiviral particle production, a packaging mixture consisting of sg*KDR*-TLCV2 vector, pMDLg/pRRE (Addgene, #12251), pRSV/REV (Addgene, #12253), and pMD2.G (Addgene, #12259) was transfected into 293T cells. MRTX1133-resistant cells were infected with lentivirus-containing medium and subsequently selected with 2 µg/mL puromycin. The resulting inducible-*KDR* KO resistant cell lines were used for further experiments. To induce *KDR* KO, cells were treated with 2 μg/mL doxycycline for 1 day; KO efficiency was confirmed by Western blot (as in Supplementary Fig. S1G).

### Deletion of the SP1 binding site on the VEGFA promoter region

To delete the predicted SP1 binding site on the VEGFA promoter, we performed site-directed mutagenesis using the Q5 Site-Directed Mutagenesis Kit (NEB Biolabs, Ipswich, MA, USA) following the manufacturer’s protocol. The deletion sequences were designed based on a prior study (Pore et al., 2004). Briefly, PCR amplification was used to generate a pUC19 vector containing or lacking the SP1 binding site. The mutated pUC19 vector was then transfected into MRTX1133-resistant cells using Fugene 4K transfection reagent (Promega, Madison, WI, USA) for further experiments.

### Enzyme-linked immunosorbent assay (ELISA)

Parental and MRTX1133-resistant cells (10,000 cells per well) were seeded in 96-well plates. After 24 hours, conditioned media (CM) were collected, and VEGFA concentration was measured using a commercial enzyme-linked immunosorbent assay (ELISA) kit (R&D Systems, Minneapolis, MN, USA) according to the manufacturer’s instructions.

### Transwell Migration Assay

We seeded 1 × 10^5^ cells in the upper chambers of Transwell inserts (Corning, NY, USA), and 500 μL of complete medium was added to the lower wells of the 24-well plate (Corning, NY, USA). After 1 day, the non-migrated cells remaining on the upper surfaces of the membrane were removed using cotton swabs. The migrated cells on the lower surface of the membrane were fixed with 4% PFA and stained with Hoechst 33342 solution (Thermo Fisher Scientific, Waltham, MA, USA). Images were acquired at 10× magnification using a fluorescence microscope (Zeiss, Göttingen, Germany) and analyzed with ZEN Blue software (Zeiss). The number of migrated cells was quantified using ImageJ software (NIH, Baltimore, MD, USA).

### Western blot

Protein lysates were extracted from cells using radioimmunoprecipitation assay (RIPA) buffer supplemented with protease and phosphatase inhibitors. Protein concentrations were determined using the bicinchoninic acid assay. Equal amounts of protein samples (10 µg) were separated on a 10% sodium dodecyl sulfate-polyacrylamide gel and transferred onto polyvinylidene difluoride membranes (Millipore, Burlington, MA, USA). The membranes were incubated overnight at 4°C with primary antibodies, followed by incubation with horseradish peroxidase (HRP)-conjugated secondary antibodies (anti-rabbit or anti-mouse, Jackson ImmunoResearch Labs, Baltimore, MD, USA) for 2 hours at room temperature. Protein expression was detected using the ChemiDoc Touch Imaging System (Bio-Rad, Hercules, CA, USA).

### Dual-luciferase reporter assay

VEGFA promoter activity was assessed using the Dual-Luciferase Reporter Assay System (Promega, Madison, WI, USA) following the manufacturer’s protocol. Briefly, parental and MRTX1133-resistant cells were seeded in 24-well plates and co-transfected with a mixture of the following plasmids using Lipofectamine 2000 (Invitrogen, Carlsbad, CA, USA): pGL4.10-VEGFprom(-1000 to -1) (Addgene, #66128), pGL4.10-VEGFprom(-1000 to -500) (Addgene, #66129), pGL4.10 empty vector (Promega, #E665A), pRL-null vector (Promega, #E2271). After 48 hours, Firefly and Renilla luciferase activities were measured using a microplate reader (BioTek Epoch, Izasa, Barcelona, Spain). Renilla luciferase values were used to normalize Firefly luciferase levels.

### Transfection of siRNA

Negative control and *SP1* siRNA were purchased from Bioneer (Daejeon, Republic of Korea). Cancer cells were transfected with a mixture of siRNA and Lipofectamine 2000 (Invitrogen, Carlsbad, CA, USA) according to the manufacturer’s instructions. After 48 hours, the transfection medium was replaced with fresh complete media, and subsequent experiments were conducted.

### Cell trafficking and tube formation

Cells were stained using CellTracker Blue and Red (Invitrogen, Carlsbad, CA, USA) according to the manufacturer’s protocol. Briefly, harvested cells were incubated in serum-free media containing 1 µM CellTracker dye for 30 minutes at 37°C. After washing with PBS, stained cells were seeded for further experiments. For the HUVEC tube formation assay, 24-well plates were coated with Matrigel Basement Membrane Matrix (Corning, NY, USA) and incubated at 37°C for 1 hour. HUVECs (1 × 10□ cells) were suspended in CM and seeded onto the Matrigel-coated wells. After 18 hours, HUVECs were stained with Calcein-AM (Corning, NY, USA) to visualize tube formation. Fluorescent images were captured using a fluorescence microscope (Zeiss) and analyzed using ZEN Blue software (Zeiss).

### Immunofluorescence

Cells and tissue samples were fixed with 4% paraformaldehyde and permeabilized with 0.2% Triton X-100. Samples were then incubated overnight at 4°C with primary antibodies, followed by Alexa Fluor 488- or 647-conjugated secondary antibodies (Invitrogen, Carlsbad, CA, USA) for 2 hours at room temperature. After washing with phosphate-buffered saline (PBS), nuclei were counterstained with a DAPI-containing mounting solution (Invitrogen). Fluorescence images were acquired using a fluorescence microscope (Zeiss) and analyzed using ZEN Blue software (Zeiss).

### Flow cytometry

Parental and MRTX1133-resistant cells were harvested, fixed with 4% paraformaldehyde, and permeabilized with 0.2% Triton X-100. Fixed cells were then incubated with anti-ZEB1 and anti-vimentin primary antibodies. After washing with PBS, cells were incubated with an Alexa Fluor 647-conjugated secondary antibody (Molecular Probes, Eugene, OR, USA). Fluorescence data were acquired using an Attune NxT Flow Cytometer (Invitrogen, Carlsbad, CA, USA) and analyzed using FlowJo software (Becton Dickinson, Franklin Lakes, NJ, USA).

### Colony-forming assay

To assess long-term proliferative capacity, 1,000 cells per well were seeded into 6-well plates and incubated for 3–5 days. The culture medium was then replaced with complete media containing doxycycline (Sigma-Aldrich, St. Louis, MO, USA) or ramucirumab (Lilly, Indianapolis, IN, USA) every 3 days for 2 weeks. At the end of the treatment period, colonies were stained with Coomassie Brilliant Blue and quantified using ImageJ software (NIH).

### Xenograft

All animal experiments were approved by the Institutional Animal Care and Use Committee (IACUC) at Seoul National University Bundang Hospital (Seongnam, Republic of Korea) under protocol No. BA-2301-360-001-04. Four-week-old female BALB/c nude mice were purchased from Koatech (Pyeongtaek, Republic of Korea) and subcutaneously injected with 1 × 10□ parental or MRTX1133-resistant PANC-1 cells mixed with Matrigel (Corning, NY, USA) and PBS. Once tumor size reached 100 mm³, mice were randomly assigned to treatment groups and received: Vehicle control, MRTX1133 (orally administered; 100 μL, daily), DC101 (intraperitoneal injection; 100 μL, once a week), and Combination therapy (MRTX1133 + DC101). Tumor size was measured using a caliper, and tumor volumes were calculated using the formula: Tumor volume = (width)^2^ × (height)/2.

### Statistical analysis

All statistical analyses were performed using Prism V9 software (GraphPad, Boston, MA, USA). Data are presented as mean ± standard deviation (SD). Comparisons between two groups were conducted using Student’s t-test. Comparisons among multiple groups were performed using one-way analysis of variance (ANOVA) followed by Tukey’s *post hoc* test for pairwise comparisons. Statistical significance was indicated as follows: *P* < 0.05 (*), *P* < 0.01 (**), *P* < 0.001 (***).

## Declarations

### Ethics approval and consent to participate

All animal experiments were approved by the Institutional Animal Care and Use Committee of the Seoul National University Bundang Hospital, Seongnam, Republic of Korea (BA-2301-360-001-04).

### Availability of data and materials

scRNA-seq and RNA-seq data have been deposited in the Gene Expression Omnibus (GEO) under accession codes GSE290526 and GSE290487, respectively. The data are available by request to the corresponding authors.

### Author contributions

**Conceived and designed the analysis:** Hwang S-H, Bae M, Kim J-W, Lee EA, and Ku J-L

**Collected the data:** Hwang S-H, Bae M, Kim J-W, Hyun SY, Lee EA, and Ku J-L

**Contributed data or analysis tools:** Hwang S-H, Bae M, Kim J-W, Hyun SY, Kim K-J, Choe JH, Kim MJ, Park JW, Jeong S-Y, Choi S, Park W, Seo J, Chae H, Kang M, Jung EH, Suh KJ, Kim SH, Kim JW, Kim YJ, Kim JH, Park H, Aguirre AJ, Lee EA, Ku J-L, and Lee K-W

**Performed the analysis:** Hwang S-H, Bae M, Kim J-W, Lee EA, and Ku J-L

**Wrote the paper:** Hwang S-H, Bae M, Kim J-W, Hyun SY, Kim K-J, Choe JH, Kim MJ, Park JW, Jeong S-Y, Choi S, Park W, Seo J, Chae H, Kang M, Jung EH, Suh KJ, Kim SH, Kim JW, Kim YJ, Kim JH, Park H, Aguirre AJ, Lee EA, Ku J-L, and Lee K-W

## Acknowledgment

None

## Supplementary Figure Legends

**Supplementary Figure S1. Angiogenesis driven by acquired resistance to MRTX1133.**

(A, B) Viability of parental and MRTX1133-resistant patient-derived organoids (PDOs) (A) cell lines (B) after treatment with MRTX1133. The experiment was repeated four times. Error bars indicate the mean□±□SD.

(C) GSVA results using MSigDB for each cluster. Each term was sorted by the difference between the average GSVA score of parental-related clusters (C1, C2, C3) and the average GSVA score of resistant-related clusters (C6, C7, C8)

(D) VEGFA concentration in P-CM and R-CM. The experiment was performed in triplicate. *P <□0.05, **P <□0.01, and ***P <□0.001.

(E) Levels of VEGFR2 and p-VEGFR2 in PANC-1-parental and resistant cells after treatment with CM derived from PANC-1-parental and resistant cells. The values were normalized by vinculin.

(F) Colony-forming assays evaluating the proliferative effect of ramucirumab (20 μg/mL for 2 weeks) on parental and resistant cells.

(G) Levels of VEGFR2 and vinculin in TLCV2-sg*KDR*-PANC-1-R cells after being treated with indicated concentrations of doxycycline (Doxy) for 1 day. The normalized values by vinculin were indicated.

(H) Colony-forming assays evaluating the proliferative effect of inducible-*KDR* KO by treatment with doxycycline (2000 ng/mL for 2 weeks) on TLCV2-sg*KDR* resistant cell lines.

(I) VEGFA concentration in CM from TLCV2-sg*KDR* resistant cells after treatment with 2000 ng/μL doxycycline. The experiment was performed in triplicate. ***P <□0.001.

(J) Resistant cells are sorted into VEGFR2-high (red) and -low (blue) populations based on the membrane VEGFR2 level.

**Supplementary Figure S2. VEGFA/VEGFR2-mediated *SP1* transcription in resistant cells.**

(A) Levels of SP1, Lamin B, and α-tubulin in fractionated nuclear and cytoplasmic protein samples of AGS and SNU-C2B parental and resistant cells. The SP1 level was normalized by the Lamin B level.

(B) Levels of SP1, Lamin B, and α-tubulin in fractionated cytoplasmic and nuclear protein samples of resistant cells following ramucirumab treatment. The SP1 level was normalized by the Lamin B level.

(C) Levels of SP1, Lamin B, and α-tubulin in fractionated cytoplasmic and nuclear protein samples of TLCV2-sg*KDR*-PANC-1-R cells after VEH or doxycycline treatment. The SP1 level was normalized by the Lamin B level.

(D) Levels of SP1, Lamin B, and α-tubulin in fractionated nuclear and cytoplasmic protein samples of PANC-1-P cells treated with 10 ng/mL of rhVEGFA. The SP1 level was normalized by the Lamin B level.

(E) Viability of parental cells treated with 10 ng/mL of rhVEGFA or vehicle and/or MRTX1133. All experiments were repeated four times. Data represent the mean□±□SD.

(F) Viability of PANC-1-R cells transfected with negative control and *SP1* siRNA after treatment with MRTX1133. All experiments were repeated four times. Data represent the mean□±□SD.

**Supplementary Figure S3. PI3Kγ activation by interaction of p110γ-p101 with KRAS reinforces the VEGFA-VEGFR2 signaling loop.**

(A) Levels of p-AKT, AKT, and vinculin in parental and resistant cells. The values were normalized by vinculin.

(B) Levels of SP1, Lamin B, and α-tubulin in fractionated cytoplasmic and nuclear protein samples of PANC-1-R cells after MK2206 treatment. The SP1 level was normalized by the Lamin B level.

(C) Levels of p-VEGFR2, VEGFR2, p-AKT, AKT, and vinculin in TLCV2-sgKDR-Resistant PANC-1 cells treated with doxycycline (2 μg/μL) and/or MRTX1133 (1 μM).

(D) The KRAS protein in AGS, SNU-C2B, and PANC-1 parental and resistant cells is immunoprecipitated, and the associated p85, p101, p110α, p110β, p110γ, KRAS, and vinculin protein levels are analyzed. The values were normalized by KRAS.

(E) Levels of p-VEGFR2, VEGFR2, p-AKT, AKT, and vinculin in resistant cells after treatment with 1 μM of eganelisib. The values were normalized by vinculin.

(F) VEGFA concentration in CM derived from resistant cells after VEH or eganelisib treatment. The experiment was performed in triplicate. **P <□0.01 and ***P <□0.001.

(G) Combination index values were calculated using CalcuSyn software according to the Chou-Talalay method after the combination treatment of eganelisib and MRTX1133 in MRTX1133-resistant cell lines.

(H) Transcriptional levels of *KRAS* and *PIK3CG* in parental and resistant cells. ***P <□0.001.

(I) Levels of KRAS, p110γ, and vinculin in PANC-1 parental and resistant cells after treatment with 10 μg/mL of CHX and 10 μM of MG132 at 1, 2, 4, and 8 hours. The values were normalized by vinculin.

**Supplementary Figure S4. CM-mediated interaction between MRTX1133-resistant cells and HUVECs.**

(A) Representative fluorescence images of HUVECs seeded on Matrigel pre-coated plates and treated with P-CM and R-CM for 18 h, then stained with Calcein AM. Scale bar: 200 μm.

(B) Representative co-localized image of HUVECs with PANC-1 parental and resistant cells by the Manders’ overlap coefficient.

(C) The number of PANC-1-R cells after treatment with complete media or CM from HUVECs for 5 days. The number of cells was automatically counted using the NYONE software (SYNENTEC GmbH). Data are presented as the mean ± SD. The experiment was independently carried out five times. **P <□0.01.

(D) GSVA results using MSigDB for P-PDO^-HUVEC^, P-PDO^+HUVEC^, R-PDO^-HUVEC^, and R-PDO^+HUVEC^. Each term was sorted by the GSVA score of P-PDO^-HUVEC^.

(E) GSVA results using MSigDB for HUVEC^-PDO^, HUVEC^+P-PDO^, HUVEC^+R-PDO^. Each term was aligned by the GSVA score of HUVEC^-PDO^.

**Supplementary Figure S5. Anti-tumor effect of VEGFR2 antibody and/or MRTX1133 in a mouse model.**

(A) PANC-1-P cell-injected mice were randomly divided into 4 groups (n = 5 in each group) and treated as indicated. The tumor volumes were recorded for 15 days.

(B) The TGI ratio of the mice bearing PANC-1-P tumor receiving MRTX1133 and/or DC101 for 15 days is calculated relative to the VEH values. Data represent the mean□±□SD (n = 5). ***P <□0.001.

(C) The body weight of mice bearing the PANC-1-R tumor receiving the indicated treatments for 15 days. Data represent the mean ±□SD (n = 5).

