## Supplementary Figures for "VEGFR2 blockade overcomes acquired KRAS G12D inhibitor resistance driven by PI3Kγ activation"

### Supplementary Figure 1

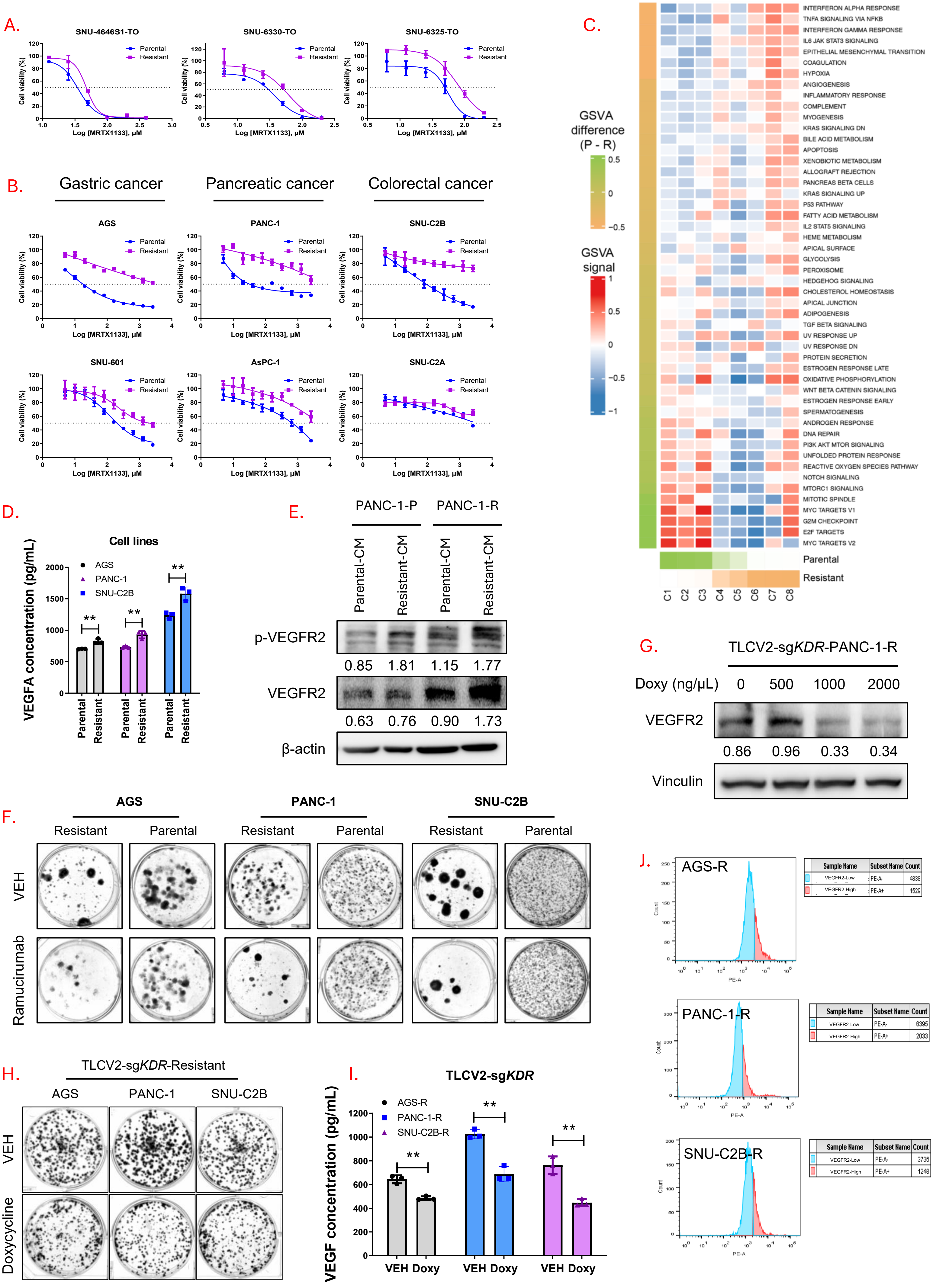

#### Supplementary Figure 2

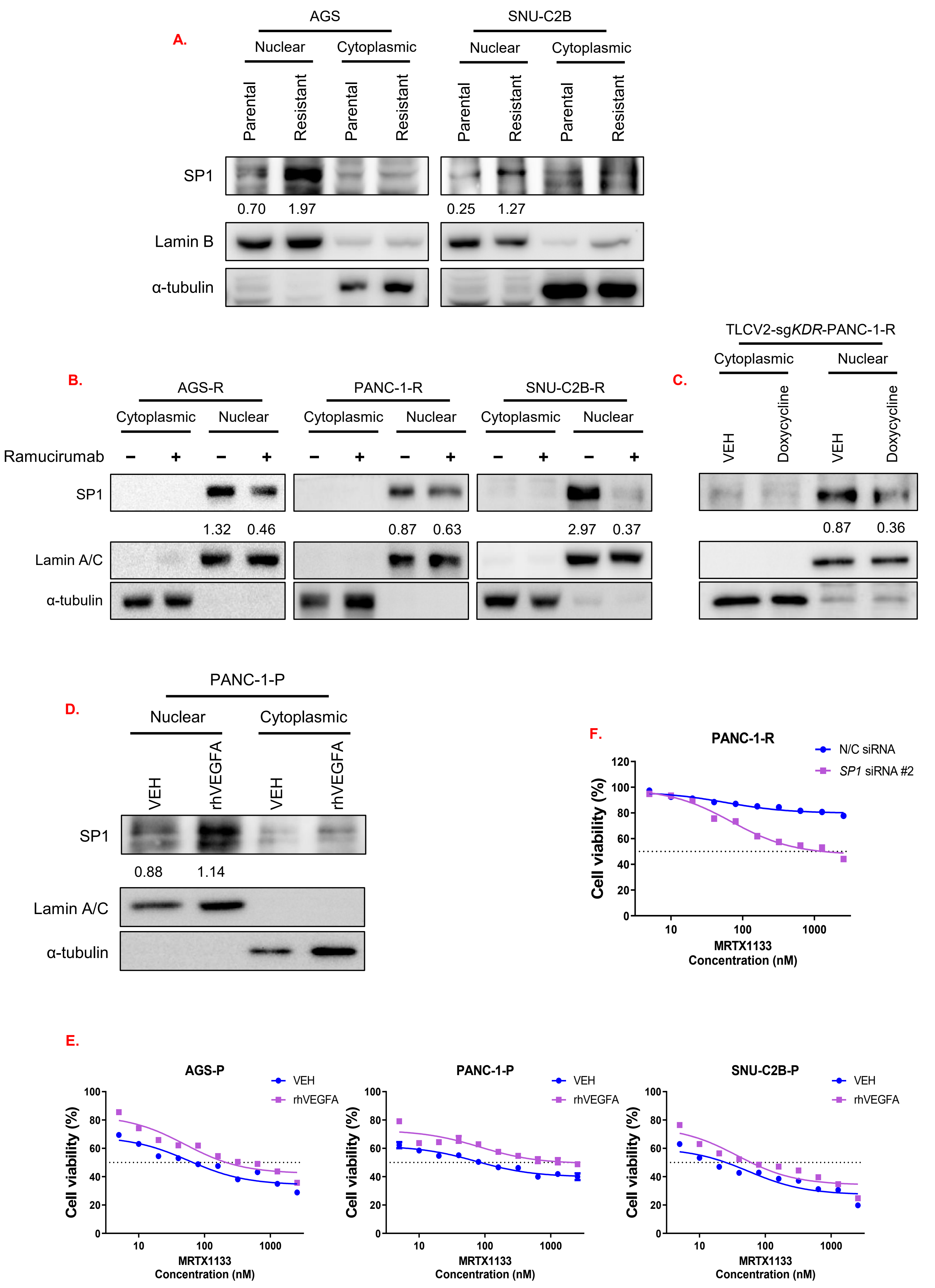

### Supplementary Figure 3

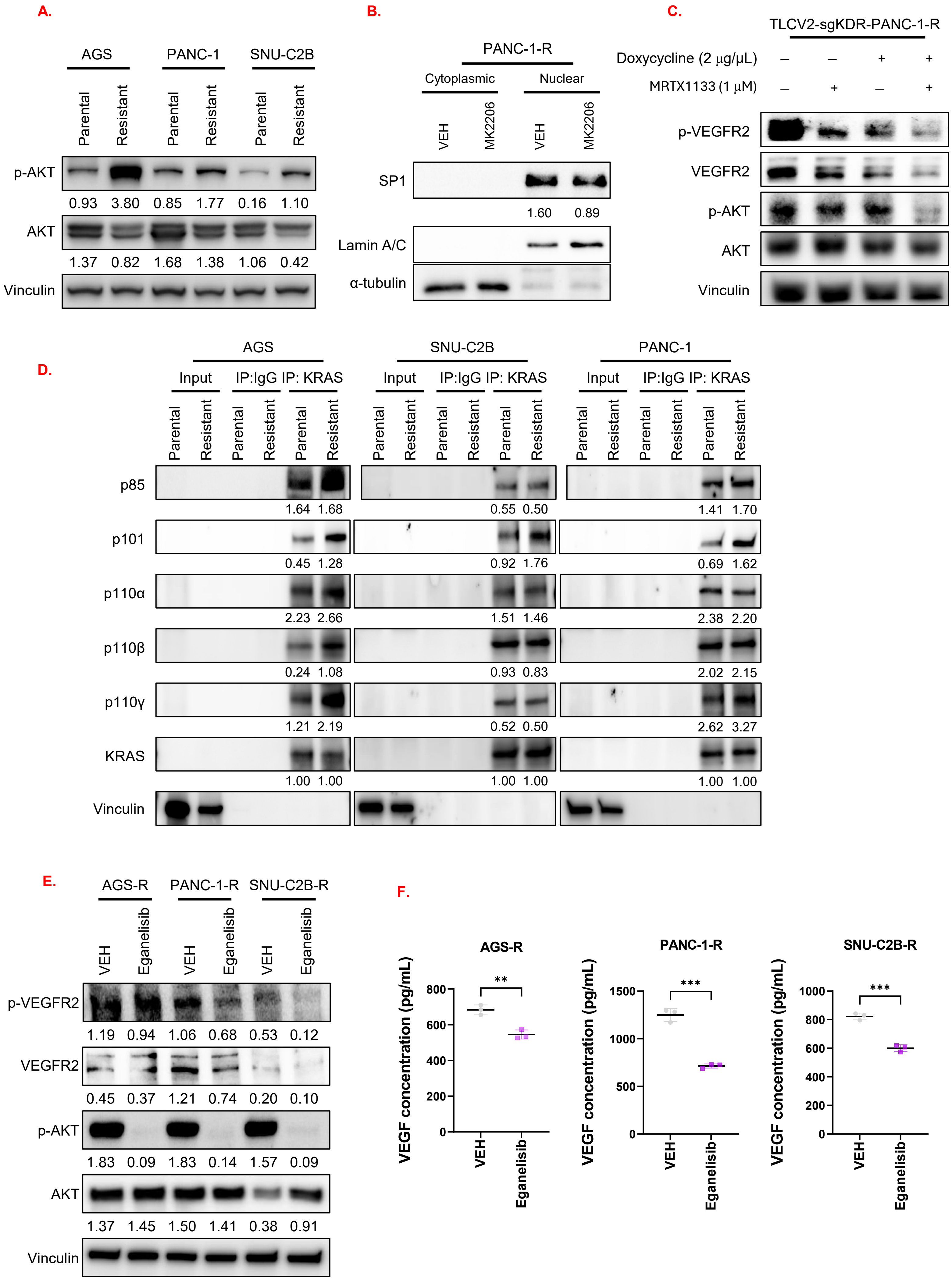

### Supplementary Figure 3

G.

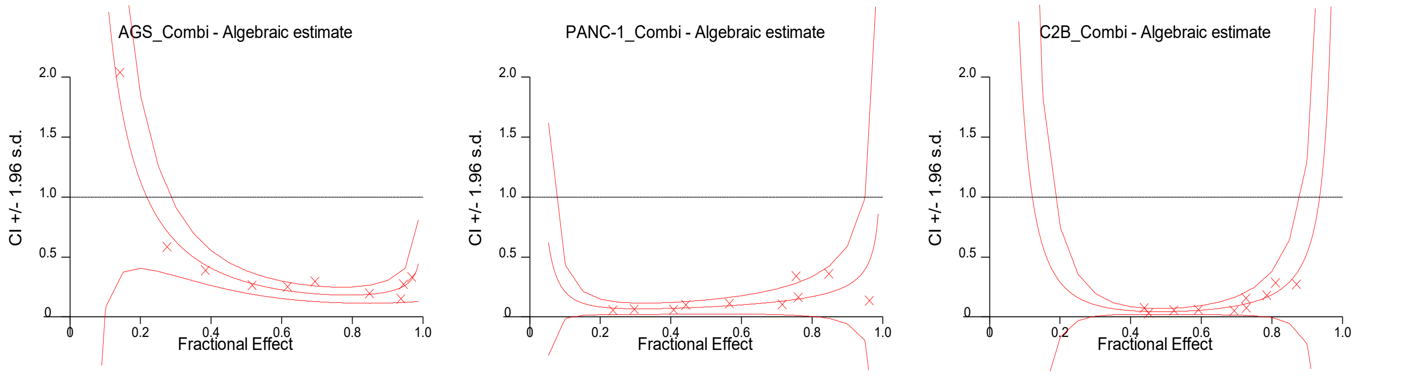


H.

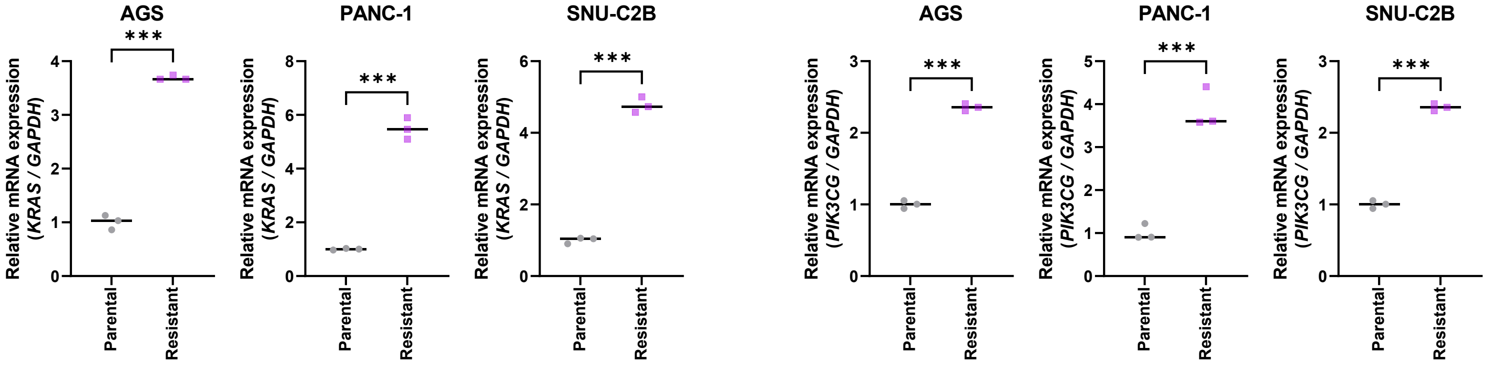


I.

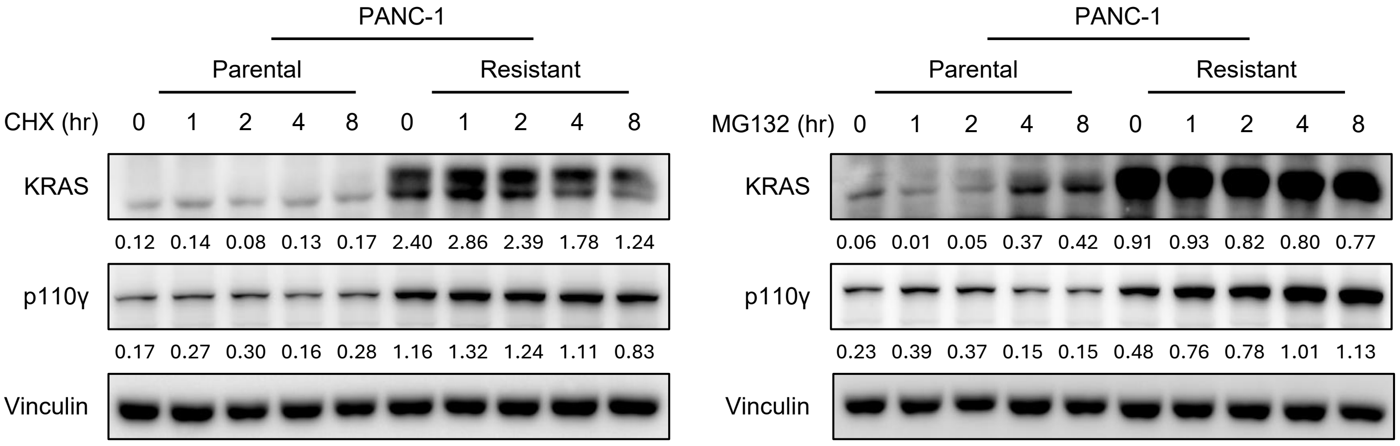

### Supplementary Figure 4

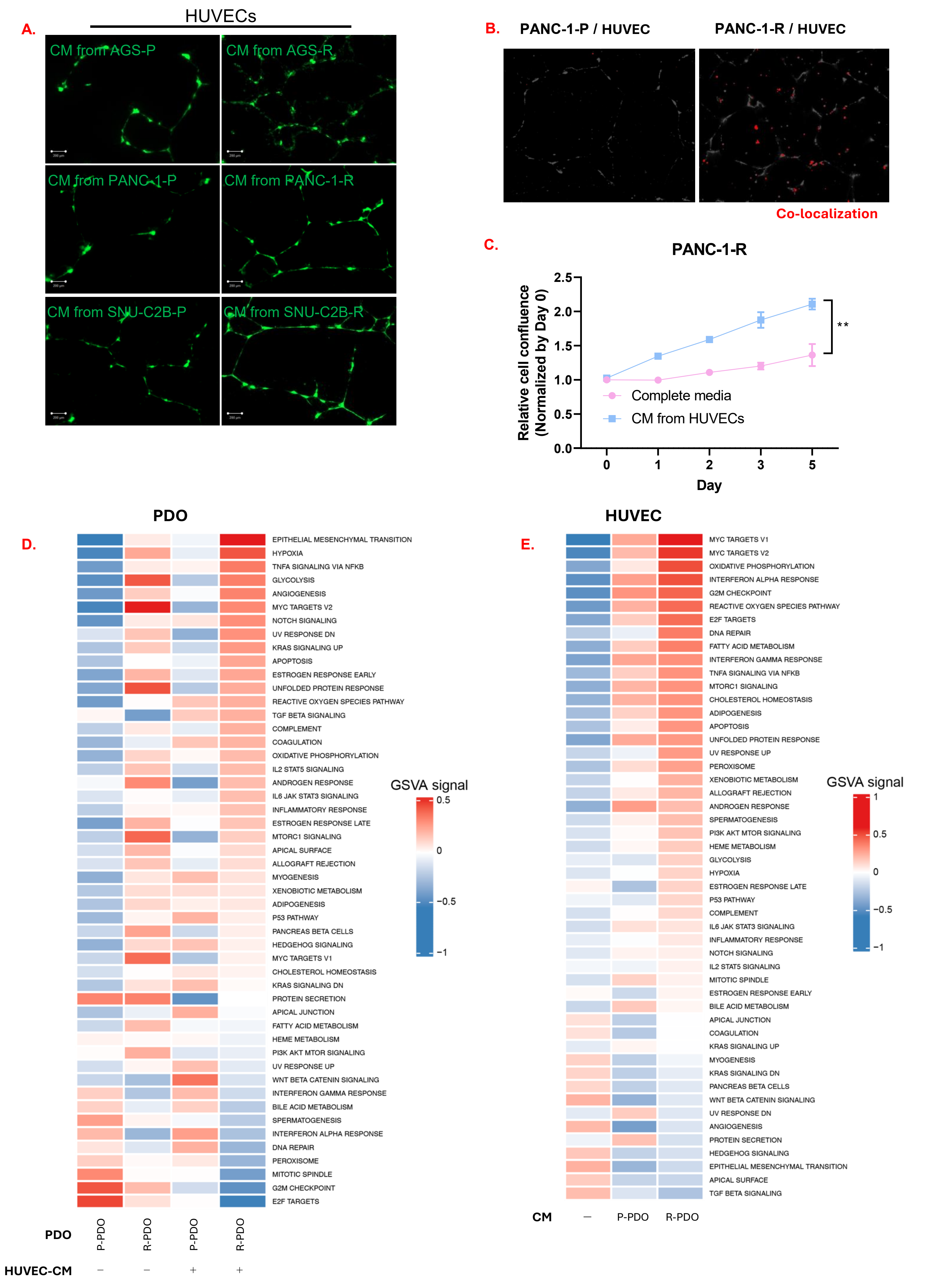

#### Supplementary Figure 5

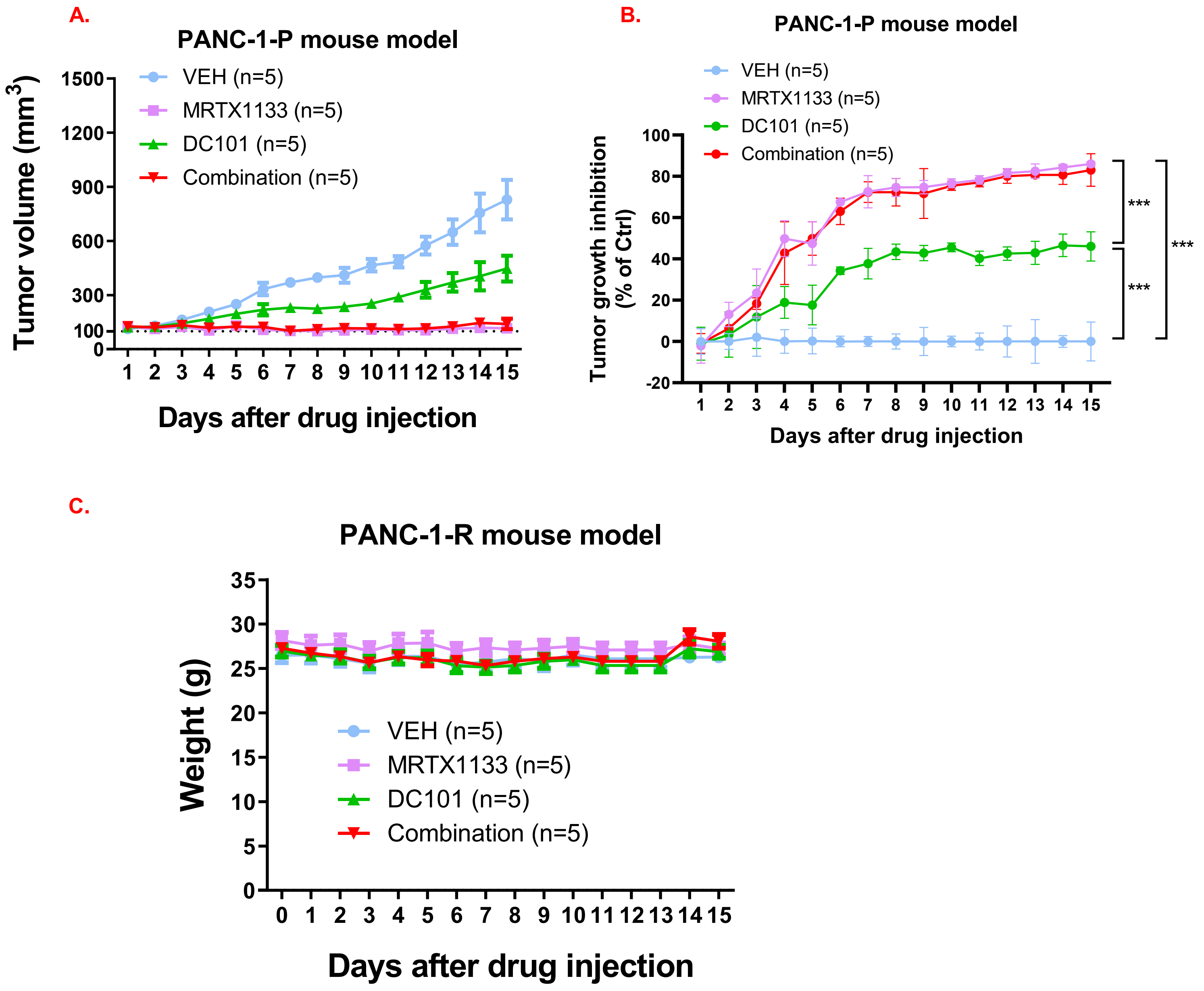
